# In vivo CRISPR base editing for treatment of Huntington’s disease

**DOI:** 10.1101/2024.07.05.602282

**Authors:** Shraddha Shirguppe, Michael Gapinske, Devyani Swami, Nicholas Gosstola, Pankaj Acharya, Angelo Miskalis, Dana Joulani, Maddie G. Szkwarek, Abhishek Bhattacharjee, Gianna Elias, Michelle Stilger, Jackson Winter, Wendy S. Woods, Daphine Anand, Colin K.W. Lim, Thomas Gaj, Pablo Perez-Pinera

**Affiliations:** Department of Bioengineering, University of Illinois Urbana-Champaign; Carl R. Woese Institute for Genomic Biology, University of Illinois Urbana-Champaign; Department of Molecular and Integrative Physiology, University of Illinois Urbana-Champaign; Carle Illinois College of Medicine, University of Illinois Urbana-Champaign; Cancer Center at Illinois, University of Illinois Urbana-Champaign

**Keywords:** Gene editing, CRISPR, Huntington’s disease, exon skipping, base editing, AAV, YAC128 mice, caspase-6, SpRY-Cas9, ABE8e, CBE4max

## Abstract

Huntington’s disease (HD) is an inherited and ultimately fatal neurodegenerative disorder caused by an expanded polyglutamine-encoding CAG repeat within exon 1 of the huntingtin (HTT) gene, which produces a mutant protein that destroys striatal and cortical neurons. Importantly, a critical event in the pathogenesis of HD is the proteolytic cleavage of the mutant HTT protein by caspase-6, which generates fragments of the N-terminal domain of the protein that form highly toxic aggregates. Given the role that proteolysis of the mutant HTT protein plays in HD, strategies for preventing this process hold potential for treating the disorder. By screening 141 CRISPR base editor variants targeting splice elements in the HTT gene, we identified platforms capable of producing HTT protein isoforms resistant to caspase-6-mediated proteolysis via editing of the splice acceptor sequence for exon 13. When delivered to the striatum of a rodent HD model, these base editors induced efficient exon skipping and decreased the formation of the N-terminal fragments, which in turn reduced HTT protein aggregation and attenuated striatal and cortical atrophy. Collectively, these results illustrate the potential for CRISPR base editing to decrease the toxicity of the mutant HTT protein for HD.

## INTRODUCTION

Huntington’s disease (HD) is an inherited and ultimately fatal neurodegenerative disorder caused by an abnormal expansion of a CAG trinucleotide repeat within exon 1 of the huntingtin (HTT) gene. This expansion leads to the production of a mutant protein that can form insoluble aggregates that contribute to the selective and progressive loss of neurons in the striatum, cerebral cortex, and other parts of the brain [1–3]. There remains no cure for HD, and current treatments offer only symptomatic relief.

Though the exact mechanism by which the mutant HTT (mHTT) protein destroys neurons remains unknown, the N-terminal domain of the mutant protein is believed to play an important role in HD. For instance, fragments of the N-terminal domain are found in nuclear inclusions in the brains of HD patients [4] while its overexpression is sufficient to induce a severe phenotype that recapitulates aspects of the disorder [5, 6].

A central event in the formation of these N-terminal fragments is the proteolysis of the mHTT protein at Asp 586 by the cysteine protease caspase-6 [7, 8]. In addition to releasing the N-terminal HTT1-586 fragment, which is subsequently processed to even smaller fragments that carry a risk for aggregation, proteolysis of the mHTT protein by caspase-6 has been hypothesized to initiate a feed-forward loop that promotes its own activation and exacerbates the pathogenesis of HD [9–11]. Importantly, it has been found that genetically ablating the proteolytic cleavage site for caspase-6 in the mHTT gene can prevent the formation of the N-terminal HTT1-586 fragment [12] and limit its subsequent proteolysis [13], which was shown to provide protection from neuronal dysfunction and neurodegeneration [12]. As such, strategies for inhibiting the formation of the N-terminal domain of the mHTT protein hold potential as therapies for HD.

One approach that can be utilized to modulate the formation N-terminal fragments of the mHTT is exon skipping. As a consensus proteolytic cleavage site for caspase-6 is fortuitously encoded in the sequence spanning exons 12 (amino acid: Ile) and 13 (amino acids: Val-Leu-Asp-Gly) of the HTT gene, technologies capable of skipping either exon could be used to produce HTT protein isoforms lacking this site [12, 14, 15] (**Fig. 1a**). This in turn is expected to prevent the proteolysis and aggregation of the mHTT protein without fully depleting the reservoir of full-length (FL) HTT protein.

**Figure 1.**
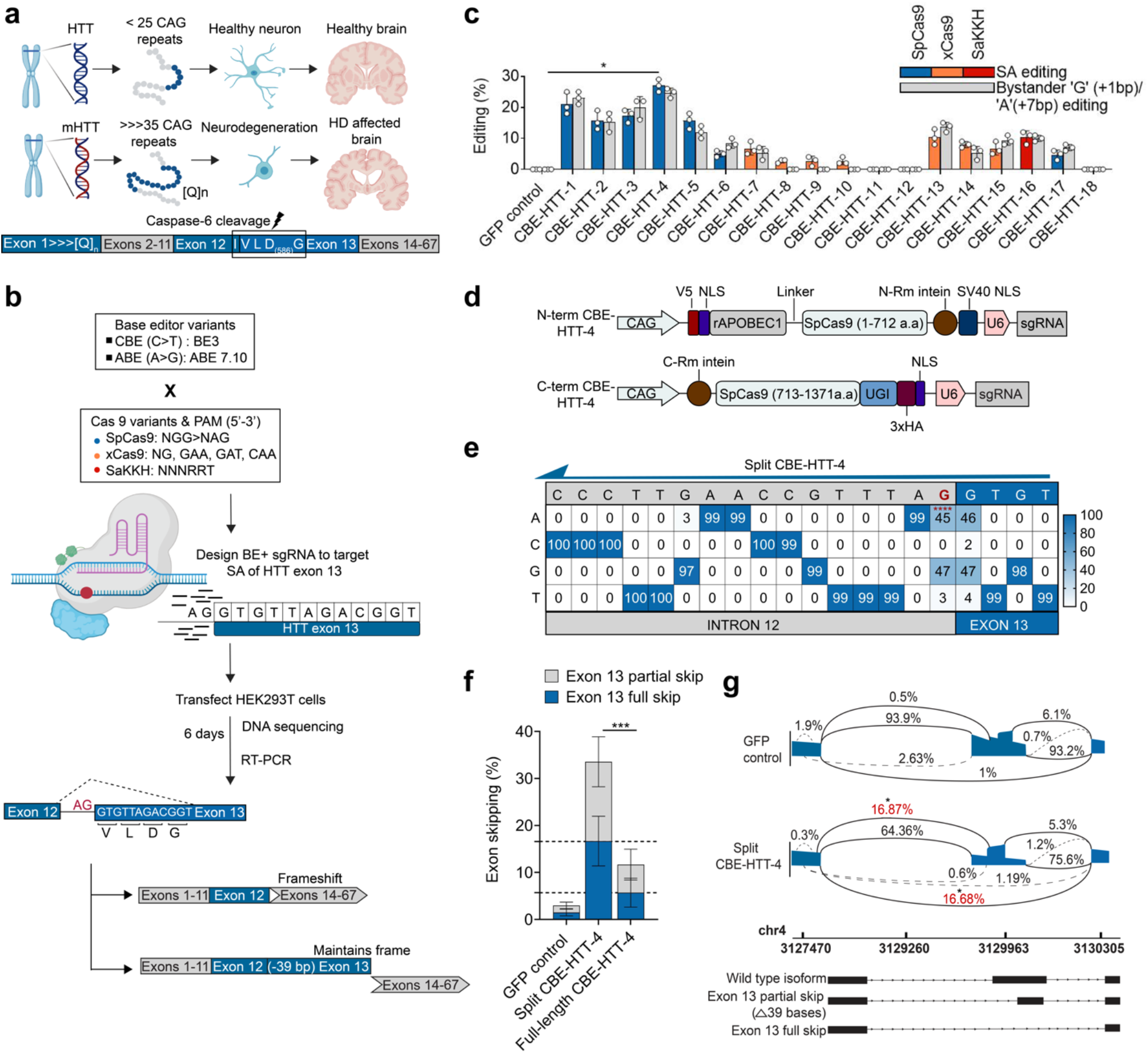
CRISPR base editing of HTT exon 13 splice acceptor (SA) sites. **(a)** (Top) Schematic representation of the pathogenesis of Huntington’s disease and (bottom) the location of the caspase-6 cleavage site in the HTT protein. **(b)** Overview of the strategy for targeting the SA for HTT exon 13 by CRISPR base editing. **(c)** Editing rates in HEK293T cells for the base editor variants targeting the exon 13 SA in HTT, as measured by next-generation sequencing (NGS; n = 3). **(d)** Schematic of the split-intein architecture used for AAV delivery. Abbreviations: ITR, inverted terminal repeat; CAG, cytomegalovirus early enhancer/chicken β- actin promoter; Rm, *Rhodothermus marinus*, NLS, nuclear localization signal; UGI, uracil glycosylase inhibitor; U6, human U6 promoter; 3xHA, three repeats of the human influenza hemagglutinin (HA) epitope; V5, epitope of the V protein of the paramyxovirus of the simian virus 5 family. **(e)** Heat map showing the editing frequencies for the split-intein version of CBE-HTT-4 in HEK293T cells, as determined by NGS (n = 3). **(f)** Quantification of exon skipping rates for the full-length and split-intein version of CBE4-HTT-4 in HEK293T cells, as determined by NGS (n = 3). **(g)** Sashimi plot depicting the splicing outcomes following treatment with CBE-HTT-4. Two major splicing patterns are shown: partial skipping of exon 13 and full skipping of exon 13. A GFP-encoding plasmid was used as a control for all experiments. All data points are biologically independent samples. Values are means and error bars indicate S.D. *P < 0.05, ***P < 0.001, ****P < 0.0001. Data for **(c)**, **(e)** and **(f)** compared using a one-tailed unpaired t-test. Data for **(g)** compared using a one-way ANOVA with Tukey’s post-hoc test.

An emergent technology capable of inducing exon skipping is CRISPR base editing. Typically consisting of a Cas9 nickase domain fused with either a cytosine deaminase for C > T editing or an adenosine deaminase for A > G editing [16, 17], CRISPR base editors are a gene-editing technology that we [18–20] and others [21–23] have shown can facilitate targeted exon skipping via their editing of highly conserved “AG” dinucleotide motifs in splice acceptor (SA) sequences. Notably, this editing can prevent the recognition of a SA by the spliceosome and promote the skipping of the targeted exon. Building on this enabling capability and reasoning that skipping of exon 12 or 13 of the HTT gene could facilitate the production of HTT protein isoforms more resistant to proteolysis and aggregation, we sought to determine the ability for CRISPR base editors to induce therapeutic exon skipping for HD.

Here, we screened 141 base editors targeting multiple splicing elements in the HTT gene, which enabled the identification of CRISPR base editing platforms that by disruption of the SA for exon 13 generate HTT protein isoforms resistant to proteolysis by caspase-6. When delivered intra-striatally to a transgenic rodent model of HD, these base editors induced skipping of HTT exon 13, an outcome that disrupted the proteolysis of the mHTT protein in vivo, leading to a substantial decrease in the accumulation of mHTT aggregates and prevention of atrophy in both the striatum and cortex. More broadly, our results illustrate the potential for base editing and exon skipping technologies to modulate the proteolysis of a target protein and demonstrate the broad potential of CRISPR base editing for treating HD.

## RESULTS

### Identification of CRISPR base editors to modify the SA of HTT exon 13

The human HTT gene consists of 67 exons, with the proteolytic cleavage sites implicated in the formation of the N-terminal fragment of the HTT protein located in exons 12 and 13 [24–26]. To disrupt the proteolysis of HTT without fully ablating its expression, we sought to use exon skipping to remove these cleavage sites, which we hypothesized would create HTT protein isoforms resistant to proteolysis. To accomplish this goal, we utilized CRISPR base editing, a gene-editing technology that can facilitate exon skipping through its targeted editing of splice sites.

First, we designed cytosine base editors (CBEs) and adenine base editors (ABEs) – initially centered on BE3 [16] and ABE 7.10 [17] respectively – to target the SAs for exons 12 and 13, as well as two putative splice enhancers for exon 12, which we identified using ESEfinder [27] and Human Splicing Finder [28]. In conjunction with these four splice sites, we also designed base editors to target the predicted caspase-3 and caspase-6 cleavage sites in exons 12 and 13 (**Fig. S1a**). To enable targeting, we utilized a suite of Cas9 variants, including the prototypical Cas9 nuclease from Streptococcus pyogenes (SpCas9), which recognizes a 5’-NGG-3’ [29] and to a lesser degree a 5’-NAG-3’ [30] protospacer adjacent motif (PAM) [16, 17] as well as the engineered variants xCas9 [31] and SaCas9-KKH [32, 33], which each recognize non-NGG PAM sequences (**Table S2**).

Of the 103 variants screened, DNA sequencing revealed that 40 modified their target sequence, with 25 of these variants found to target exon 12, including its SA, its splice enhancers and the predicted protease cleavage sites (**Fig. S1b**). However, only two of these 25 variants edited their targets with an efficiency over 20% (**Fig. S1b**), with a further analysis revealing that neither of these two variants induced skipping at a rate that exceeded 10% (**Fig. S1c**). Notably, we identified 15 variants that edited their respective target sequence at the SA of HTT exon 13, with one variant, CBE-HTT-4, installing edits with 27% efficiency (P = 0.0001; **Fig. 1C**).

We next compared the DNA editing capabilities of the most efficient CBE system identified above, CBE-HTT-4, against a split-intein version of it intended for AAV delivery [19, 34–37], as the carrying capacity of an AAV vector limits its ability to deliver full-length versions of these editors within a single viral particle [38]. Our split-intein CBE is encoded across two vectors: one vector expresses the APOBEC deaminase and the first 712 residues of the SpCas9 protein fused to the N-terminus DnaB intein from *Rhodothermus marinus,* which mediates a protein *trans*- splicing reaction with its C-terminal counterpart, and the second vector expresses the C-terminal DnaB intein fused to residues 713-1,371 of the SpCas9 protein fused with an uracil glycosylase inhibitor (UGI) domain. Each vector further encodes an sgRNA expression cassette (**Fig. 1D**).

This split-intein system introduced the target C > T edit in the SA of HTT exon 13 at an efficiency of ∼45% in HEK293T cells, though we found that it also created a G > A bystander mutation +1-bp from the SA at an efficiency of ∼46%, with this edit expected to create a missense mutation in HTT [GTG > ATG: Val584 > Met] (**Fig. 1e and Fig. S2**). Nonetheless, we note that exonic mutations that occur simultaneously with the target modification in the SA are not expected to have a biological effect, as the exon would be skipped.

Interestingly, ablating the SA for HTT exon 13 is predicted to result in a frameshift mutation in the HTT transcript, which is expected to trigger nonsense mediated decay (NMD) and a reduction in HTT expression. However, according to SpliceAI [39], MaxEntScan [40] and Human Splicing Finder [28, 41], there exists an alternative SA in exon 13, located 39 bps from the canonical SA, whose utilization is predicted to produce a 13-residue truncation in the HTT protein that would disrupt the caspase-6 cleavage site but not alter its reading frame (**Fig. S3a and S3b**).

We next quantified the rate of exon skipping induced by this split-intein CBE in HEK293T cells. Based on an NGS analysis of PCR amplicons generated from cDNA using primers that targeted HTT exons 11 and 14 (**Fig. S2b**), the split-intein version of CBE-HTT-4 induced more than three-fold higher skipping of exon 13 compared to its FL equivalent (∼33% vs ∼11.6%; P = 0.0006; **Fig. 1f**). Critically, this analysis further revealed that editing of the SA in HTT exon 13 did in fact generate two major products, with ∼16.7% of the analyzed HTT transcripts found to lack the entirety of exon 13, and ∼16.9% of the transcripts found to lack only the first 39-bps of exon 13 (P < 0.05 for both; **Fig. 1g**).

Last, we determined the specificity of this split-intein CBE. Using NGS, we measured C > T editing at the eight highest scoring off-target (OT) sites, as predicted by Cas-OFFinder [30]. No editing was observed in HEK293T cells at any of the candidate OT sites (P > 0.05; **Fig. S4**).

### Editing the HTT exon 13 SA site alters HTT splicing

To further characterize the expression patterns resulting from the disruption of the SA site for HTT exon 13, we generated clonal HEK293T cell lines edited by the split-intein version of CBE-HTT-4 (**Fig. 2a**). After confirming the presence of the target edit in each clone (**Fig. 2b**), we conducted an NGS analysis on PCR products from cDNA prepared from the base editor-modified cell lines, as well as naive unedited HEK293T cells. In total, ∼37% of the HTT transcripts in the modified clones lacked exon 13 (P = 0.001; **Fig. 2c**), while ∼46% of transcripts lacked only the first 39-bps of it (P = 0.0003; **Fig. 2c**). Interestingly, ∼10% of all transcripts retained a fragment of intron 12, which we hypothesized occurred through the utilization of an ‘AG’ sequence in an intergenic region 18-bp upstream of the targeted SA site (**Fig. 2d**), which is consistent with the model generated by Splice AI that predicted that disrupting the exon 13 SA and the concurrent introduction of a G > A mutation +1-bp of the SA would create cryptic SAs at nearby exonic and intronic sequences (**Fig. S3a)** .

**Figure 2.**
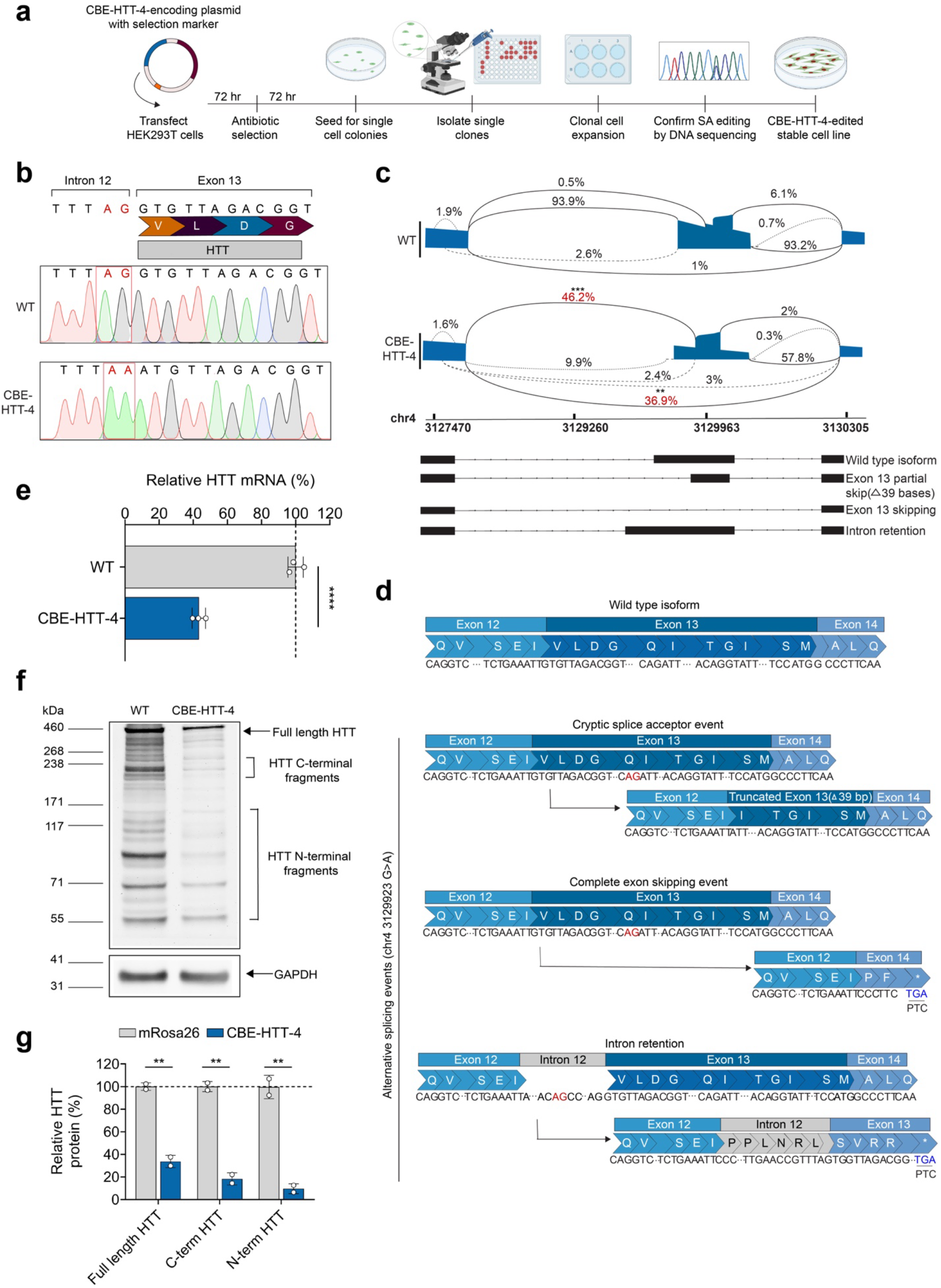
CRISPR base editing of HTT exon 13 influences splicing and protein expression. **(a)** Overview of the workflow for creating isogenic HEK293T cell lines. **(b)** (Top) Representative Sanger sequencing traces of a wild-type (WT) and (bottom) CBE-HTT-4-modified cell line with the target edit at the exon 13 SA in HTT. **(c)** Sashimi plot depicting the predicted splicing outcomes in WT cells and CBE-HTT-4-modified cells**. (d)** Representation of alternative splicing before and after utilization of a cryptic SA in HTT exon 13. Outcomes include partial skipping of exon 13, complete skipping of exon 13, and retention of the intron upstream of exon 13. **(e)** Relative HTT mRNA in WT and CBE-HTT-4-modified cells (n = 3). **(f)** Western blot of HTT protein from WT and CBE-HTT-4-modified cells following incubation with purified caspase-6 enzyme. **(g)** Quantification of relative full-length HTT protein, C-terminal and N-terminal HTT protein fragments in the WT or CBE-HTT-4-modified cells. All values are normalized to GAPDH (n = 3). Values are means and error bars indicate S.D. **P < 0.01, ***P < 0.001, ****P < 0.0001. Data for **(c)**compared using a one-way ANOVA with Tukey’s post-hoc test. Data for **(e)** and **(g)** and compared using a one-tailed, unpaired t-test.

As the skipping of exon 13 and the retention of intron 12 is expected to result in a frameshift mutation in HTT, which in turn is anticipated to decrease its expression by NMD, we also measured the relative abundance of HTT mRNA in the isolated clones. Based on qPCR, we found that the base editor-modified cell lines had on average a ∼57% decrease in HTT mRNA compared to the naive, unedited HEK293T cells (P < 0.0001; **Fig. 2e**).

To determine whether the HTT protein isoforms produced in the modified cell lines were resistant to proteolysis by caspase-6, we incubated their lysates with purified caspase-6 enzyme (**Fig. 2f, g**). According to western blot, the treated lysates originating from the base editor-modified clones showed a ∼67% decrease in FL HTT protein compared to the naive cells (P = 0.002, **Fig. 2g**). Importantly, we observed ∼90% (P = 0.0038, **Fig. 2g**) and a ∼82% (P = 0.001, **Fig. 2g**) decrease in N- and C-terminal HTT protein fragments [42–44], respectively, compared to their naive counterparts, indicating reduced proteolysis of HTT protein by caspase-6.

Thus, our results demonstrate that disrupting the SA for HTT exon 13 by CRISPR base editing can create HTT protein isoforms resistant to cleavage by caspase-6.

### In vivo exon skipping of HTT improves HD-related deficits

We next evaluated the effectiveness of the split-intein CBE in a transgenic rodent model of HD, specifically the YAC128 mouse model, which expresses a mutant form of the FL human HTT gene with intervening intronic sequences. YAC128 mice recapitulate pathological hallmarks of HD, including the formation of mHTT inclusions and brain atrophy [45–47].

To mediate the delivery of the HTT-targeting split-intein CBE, we used AAV9, which can transduce neurons in the striatum [48] and has demonstrated effectiveness in various studies [49–52]. For this experiment, we bilaterally injected the striatum of two-month-old YAC128 mice with 1 x 10^9^ vector genomes (VGs) total, per hemisphere, of two AAV9 vectors encoding the split-intein version of CBE-HTT-4, with the N- and C-terminal CBE-encoding AAV vectors injected at a 1:1 ratio (**Fig. 3a**). As negative control, we injected YAC128 mice with an AAV formulation encoding split-intein CBEs targeting the mRosa26 locus [53, 54], a safe harbor region in the mouse genome.

**Figure 3.**
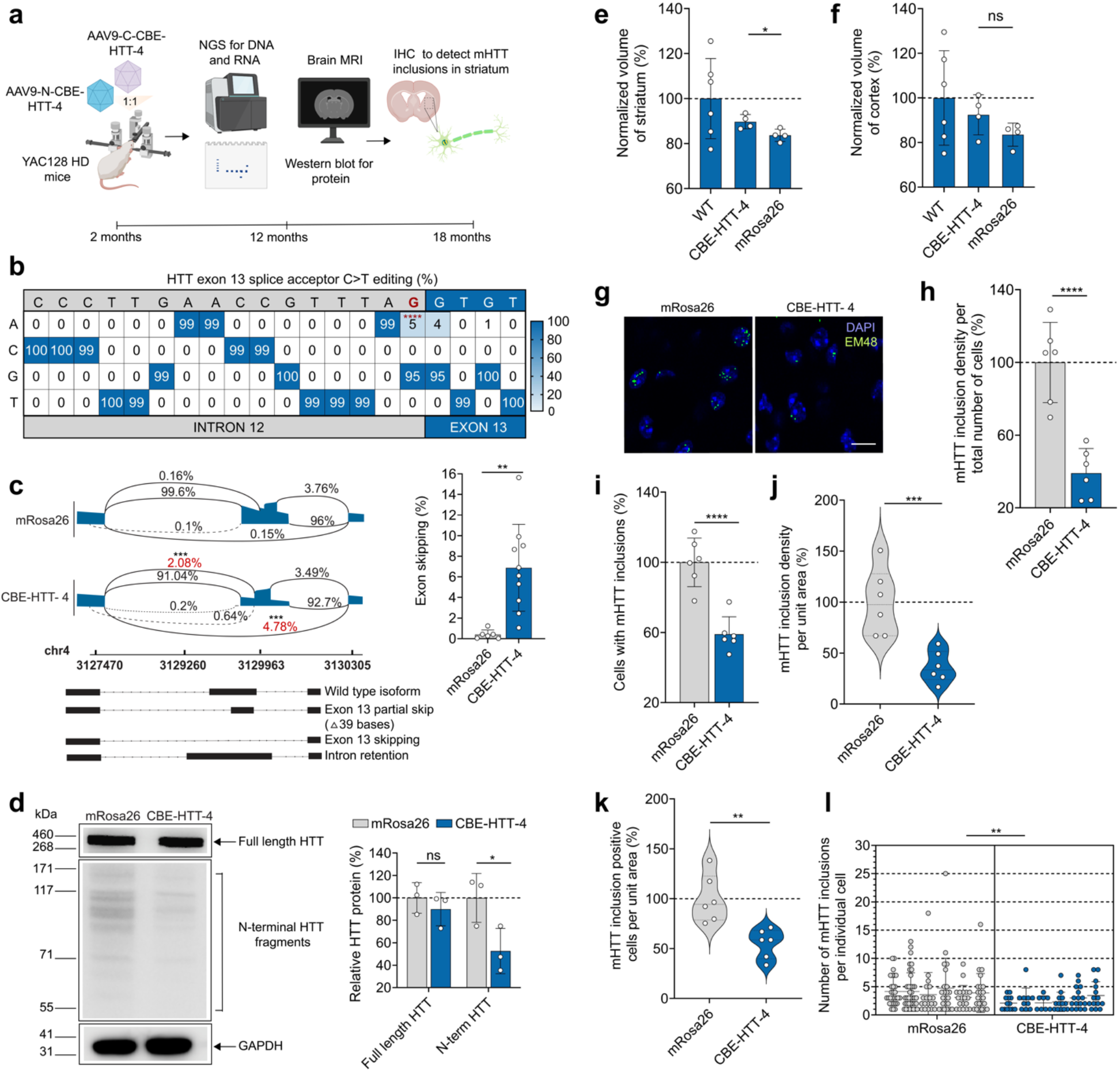
Base editing of the HTT exon 13 SA in YAC128 mice. **(a)** Schematic of the study in YAC128 mice. **(b)** DNA editing frequencies in the striatum of 12-month-old YAC128 mice injected with 1 x 10^9^ vector genomes (VGs) of AAV9-CAG-CBE-HTT-4 (n = 10) or with AAV9-CAG-CBE- mRosa26 (n = 7). **(c)** (Left) Sashimi plot depicting the splicing events in the striatum of 12-month- old YAC128 mice injected with 1 x 10^9^ VGs of AAV9-CAG-CBE-HTT-4 (n = 10) or AAV9-CAG- CBE-mRosa26 (n = 6). (Right) Summary of HTT exon 13 skipping rates, as determined by NGS. **(d)** (Left) Western blot for human HTT protein products from the striatum of 12-month-old YAC128 mice injected with 1 x 10^9^ VGs of AAV9-CAG-CBE-HTT-4 (n = 3) or AAV9-CAG-CBE-mRosa26 (n = 3). (Right) Quantification of relative protein expression normalized to GAPDH (n = 3). **(e, f)** Normalized **(e)** striatum and **(f)** cortex volumes of 12-month-old YAC128 mice injected with 1 x 10^9^ VGs of AAV9-CAG-CBE-HTT-4 (n = 4) or AAV9-CAG-CBE-mRosa26 (n = 4). All data normalized to 12-month-old wild-type FVB/NJ mice (n = 6). **(g)** Representative immunofluorescence staining of mHTT inclusions in the striatum of 12-month-old YAC128 mice injected with 1 x 10^9^ VGs of AAV9-CAG-CBE-HTT-4 (n = 6) or AAV9-CAG-CBE-mRosa26 (n = 6). Scale bar, 52 µm. **(h, i, j, k, l)** Quantification of **(h)** mHTT inclusion density per total number of cells in the striatum, **(i)** the percentage of cells with ≥1 mHTT inclusion per total number of cells, **(j)** mHTT inclusion density per unit area, **(k)** the percentage of cells with ≥1 mHTT inclusion per area unit, and **(l)** the number of mHTT inclusions per cell All data normalized to YAC182 mice injected with AAV9-CAG-CBE-mRosa26. Values are means and error bars indicate S.D. *P < 0.05, **P < 0.01, ***P < 0.001, ****P < 0.0001. Data for **(b)**, **(d, e)**, and **(h, i, j, k, l)** compared using a one-tailed unpaired t-test. Data for **(c)** compared using a one-way ANOVA with Tukey’s post- hoc analysis.

At four weeks post-injection, we conducted an immunofluorescence-based analysis to determine the expression of the CBE using an antibody that recognizes a V5 epitope contained on its N-terminus, which revealed the CBE was predominantly expressed within the striatum and the cortex (**Fig. S5**). Using NGS, we next measured DNA editing in bulk striatal tissue from 12- month-old YAC128 mice, observing the target C > T modification in the SA for exon 13 in ∼5% of the reads analyzed from mice injected with the HTT-targeting CBE (P < 0.0001; **Fig. 3b**). The previously observed G > A bystander mutation, located +1-bp of the target SA, was also found in∼4.8% of the analyzed reads (P < 0.0001; **Fig. 3b**). Notably, no editing was observed in tissue from YAC128 mice injected with AAV encoding the mRosa26-targeting CBE.

To determine if modifying the exon 13 SA influenced HTT mRNA splicing, we conducted an NGS analysis on RT-PCR products from mRNA isolated from bulk striatal tissue of 12-month-old YAC128 mice. From this analysis, we found that ∼4.8% of all HTT transcripts from mice treated with the HTT-targeting CBE lacked exon 13 (P = 0.001) and that ∼2% of all transcripts lacked the first 39-bps of exon 13 (P = 0.001; **Fig. 3c**). Importantly, no skipping was observed in samples from mice injected with the mRosa26-targeting CBE.

Using western blot, we next measured the abundance of the HTT protein in 12-month-old YAC128 mice with an anti-HTT antibody clone (5HU-1H6) that recognizes the N-terminal domain (amino acids 115-129) of the human protein [55, 56]. Within the striatum, we found that YAC128 mice treated by the split-intein CBE had ∼48% less N-terminal HTT protein (P = 0.0254; **Fig. 3d**) and 10% less FL HTT protein (P = 0.2; **Fig. 3d**) compared to animals injected with the mRosa26- targeting CBE.

We next determined if disrupting the SA for exon 13 improved HD-related deficits in YAC128. Using magnetic resonance imaging (MRI), we measured volumetric changes of the striatum and cortex of 12-month-old YAC128 mice. As expected, YAC128 mice injected with the mRosa26-targeting CBE had reduced striatal volume compared to their wild-type littermates (**Fig. 3e**); however, in YAC128 mice treated by base editing the striatum was ∼10% larger than in mice injected with the mRosa26-targeting CBE (P = 0.04; **Fig. 3e**). No preseveration in cortical volumes were observed for the mice treated by base editing (P > 0.05; **Fig. 3f**).

A hallmark of HD is the accumulation of intraneuronal inclusions consisting of the mHTT protein, which can consist of N-terminal fragments of the mHTT protein [57–61]. We thus conducted an immunofluoresence-based analysis to determine if disrupting the exon 13 SA also reduced the abundance of mHTT-containing inclusions in 18-month-old YAC128 mice. Notably, at this time point, mHTT inclusions in YAC128 mice cluster predominately in the nucleus of cells [45, 62, 63].

Using an anti-HTT antibody (EM48) that recognizes the N-terminal domain of the protein (amino acids 1–256) and is selective for mHTT-containing inclusions [64] (**Fig. 3g**), we determined the relative density of mHTT inclusions in the striatum, a ratio that we defined as the number of mHTT inclusions per the total number of DAPI^+^ cells per section. From this analysis, we measured that mice treated by base editing had a ∼61% decrease in inclusion density compared to mice injected with the mRosa26-targeting CBE (P < 0.0001; **Fig. 3h**). We further measured that mice treated by base editing had a ∼41% decrease in the percentage of cells with detectable mHTT inclusions (p < 0.0001; **Fig. 3i**), a ∼63% decrease in mHTT inclusion density per area unit (P = 0.0008; **Fig. 3j**) and a ∼45% decrease and the number of cells with detectable mHTT inclusions per area unit (P = 0.0015; **Fig. 3k**). Additionally, for those cells with detectable mHTT inclusions, we measured a ∼50% decrease in their total number of inclusions per cell (P = 0.0027; **Fig. 3l**).

Last, we evaluated if disrupting the SA for exon 13 influenced astrogliosis and microgliosis, two neuroinflammatory responses that are suspected to play a role in HD [65, 66]. Within the striatum, YAC128 mice treated by the CBE had a ∼42% decrease in the number of cells positive for the reactive astrocyte marker glial fibrillary acidic protein (GFAP) [67] compared to the controls (P = 0.03; **Fig. S6b)**. However, no difference was observed in the number of GFAP^+^ cells in the cortex for animals treated by base editing compared to controls (**Fig. S6c**). Similarily, we observed no difference between either group in the number of cells positive for the reactive microglia marker ionized calcium-binding adaptor molecule 1 (Iba-1) [68] in either the striatum or cortex (**Fig. S6e, f**).

Collectively, these findings demonstrate that CRISPR base editors can be delivered in vivo to disrupt the SA for HTT exon 13 and that their action can attenuate the formation of mHTT inclusions and preserve the striatum of YAC128 mice.

### Near-PAMless base editors enhance editing of the SA for HTT exon 13

While an early-generation SpCas9-based CBE enabled editing of the SA site for HTT exon 13, the emergence of a large catalog of base editor variants with improved efficiency, increased precision, and greater targeting range prompted us to examine whether their utilization could enhance the skipping of HTT exon 13.

To determine this, we designed 19 new CBEs comprised of the deaminase domain from CBE4max [69] and 19 new ABEs based on ABE8e [70]. To enable a more systemic tiling of the SA for HTT exon 13, we also utilized the near PAM-less Cas9 variant SpRY [71] for DNA targeting (**Fig. 4a**).

**Figure 4.**
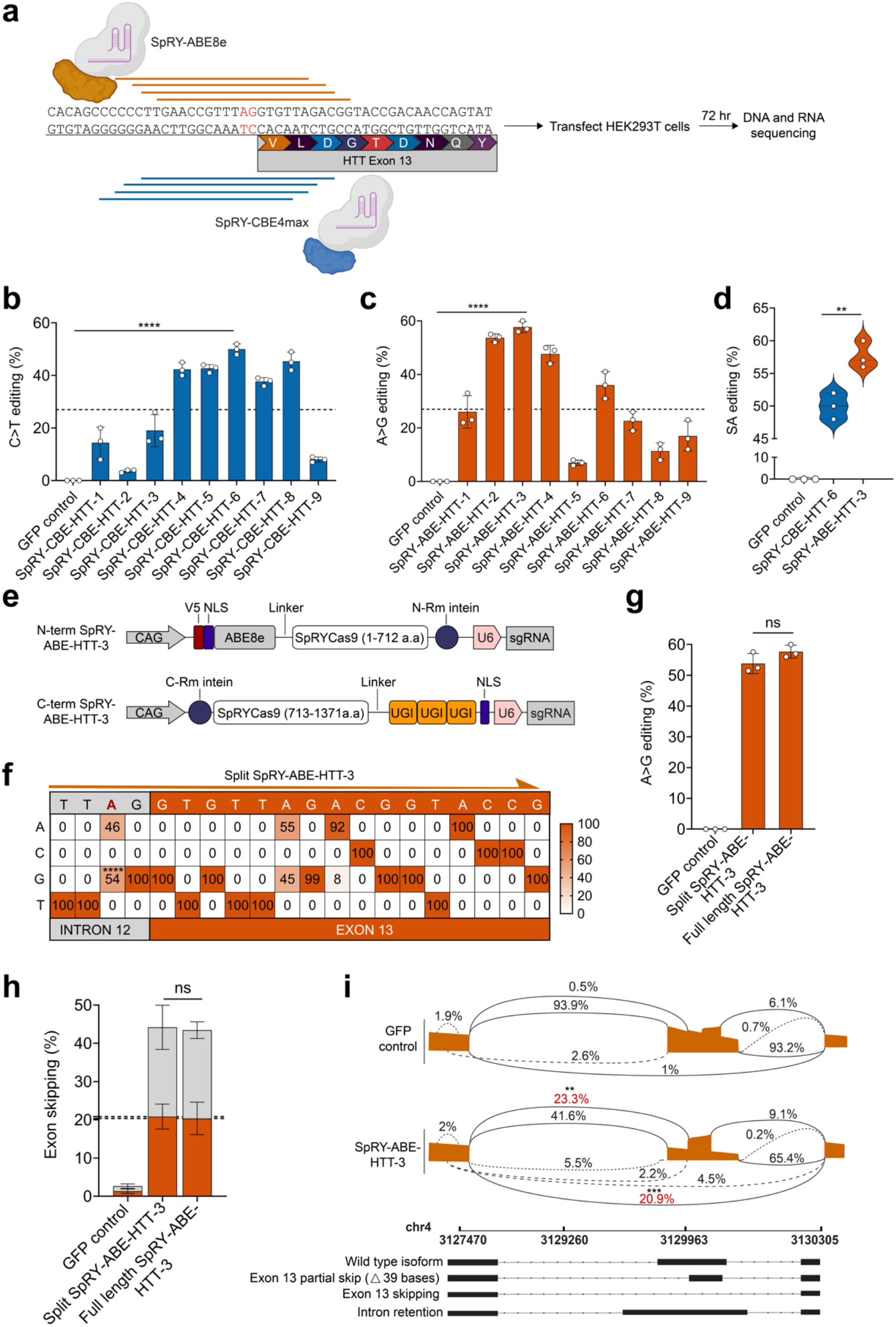
Disruption of the HTT exon 13 SA using near-PAMless CRISPR base editors. **(a)** Schematic of the approach to disrupt HTT exon 13 SA using near-PAMless base editor variants. **(b, c)** Editing rates in HEK293T cells at the exon 13 SA, as determined by NGS (n = 3). **(d)** Violin plot depicting the editing efficiencies of SpRY-CBE-HTT-6 and SpRY-ABE-HTT-3 (n = 3). **(e)** Schematic of the split-intein architecture used for AAV delivery. **(f)** Heat map depicting the editing frequencies for the split-intein version of SpRY-ABE-HTT-3 in HEK293T cells, as determined by NGS (n = 2). **(g)** Quantification of exon skipping rates induced by the full-length and split-intein version of SpRY-ABE-HTT-3 in HEK293T cells, as determined by NGS. **(h)** Quantification of the percentage of transcript variants with exon 13 partially skipped (grey) or exon 13 fully skipped (orange), as measured by NGS (n = 2). **(i)** Sashimi plot depicting the splicing events for SpRY-ABE-HTT-3: partial skipping of exon 13 or full skipping of exon 13. A GFP-encoding plasmid was used as a control for all experiments. All data points are biologically independent samples. Values are means and error bars indicate S.D. **P < 0.01, ***P < 0.001, ****P < 0.0001; Data for **(b, c, d)** and **(f, g, h)** compared using a one-tailed unpaired t-test. Data for **(i)** compared using a one-way ANOVA with Tukey’s post-hoc analysis.

Using NGS, we evaluated the editing capabilities of these 38 variants in HEK293T cells, finding that nine of the 38 editors modified the SA for HTT exon 13 with efficiencies that exceeded those for the SpCas9-based CBE variant used in our initial study (**Fig. 4b, c**). This included SpRY- CBE-HTT-6 and SpRY-ABE-HTT-3, two variants that edited the exon 13 SA with efficiencies of ∼50% and ∼57%, respectively (P < 0.0001 for both; **Fig. 4d**), rates that corresponded to a nearly two-fold improvement to the originally identified SpCas9-based CBE variant.

Following this screen, we next determined the editing capabilities of the split-intein version of SpRY-ABE-HTT-3 (**Fig. 4e**), which we utilized for all subsequent studies given its increased efficiency compared to SpRY-CBE-HTT-6. According to an NGS analysis of editing outcomes in HEK293T cells, the split-intein equivalent of this ABE induced the target A > G edit in the SA site for HTT exon 13 at an efficiency of ∼54% (P < 0.0001; **Fig. 4f**), a number that was largely indistinguishable from its FL counterpart, though both variants were found to edit a bystander ‘A’ +7-bps downstream of the SA at an efficiency of ∼45% (**Fig. 4f, S7)**. Fortuitously, this edit was expected to create only a silent mutation in HTT [TTA > TTG: Leu585 > Leu]. Additionally, the split SpRY-ABE-HTT-3 installed another bystander mutation +9-bps downstream of the SA in 8% of the reads (**Fig. 4f**), which causes a missense mutation in residue D586 [GAC> GGC: Asp586>Gly].

Using NGS, we next measured the rate of exon skipping for each ABE via the RT-PCR products from mRNA extracted from transfected HEK293T cells. We measured no difference in exon skipping for the split-intein ABE versus its full-length equivalent (P > 0.05), with both variants found to induce ∼43% skipping (**Fig. 4g**). In the case of the split-intein ABE, ∼47% of these skipping events led to the removal of the entirety of exon 13, while ∼53% of them involving the skipping of just the first 39-bps of exon 13 (P > 0.05 for both events; **Fig. 4h, i**).

Last, we analyzed OT editing by the ABE in HEK293T cells, measuring no A > G editing by NGS at any of the top five OT sites, as predicted by Cas-OFFinder [72] (P > 0.05; **Fig. S8**).

### Near-PAMless base editors edit HTT exon 13 SA in vivo and improve HD-related deficits

Given its improved editing capabilities compared to the SpCas9-based CBE used in our initial study, we next evaluated the ability for SpRY-ABE-HTT-3 to edit the SA for HTT exon 13 in YAC128 mice (**Fig. 5a**). AAV9 vectors encoding the split-intein version of this editor were bilaterally injected to the striatum of one-month-old YAC128 mice at three doses: 1 x 10^9^, 1 x 10^10^, and 1 x 10^11^ VGs total per hemisphere, with the N- and C-terminal AAV vectors injected at a 1:1 ratio.

**Figure 5.**
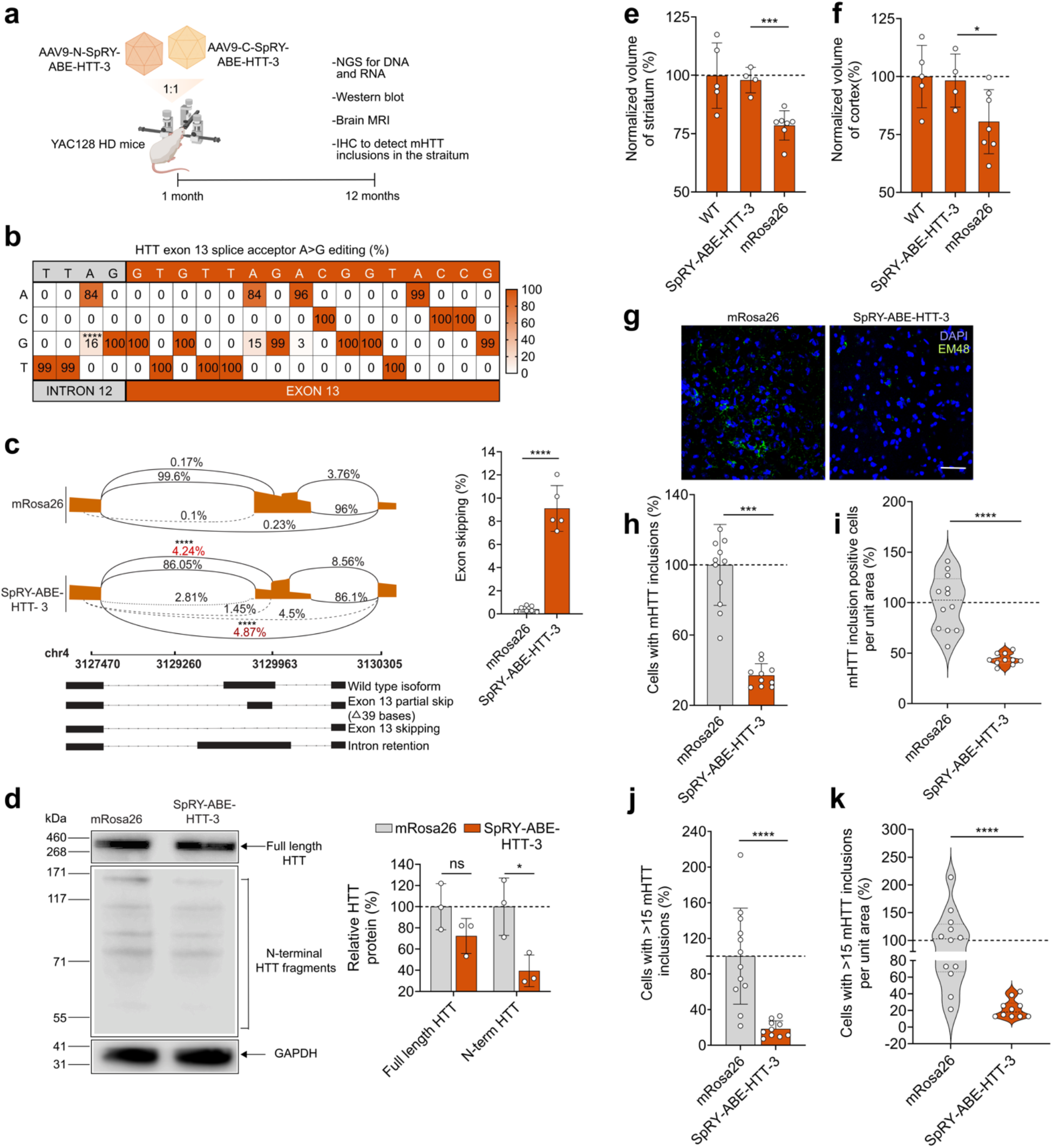
Disrupting HTT exon 13 SA using a near-PAMless base editor improves HD- related deficits in YAC128 mice. **(a)** Schematic of the experiments conducted in YAC128 mice. **(b)** DNA editing frequencies in the striatum of 12-month-old YAC128 mice injected with 1 x 10^11^ VGs of AAV9-CAG-SpRY-ABE-HTT-3 (n = 5), with a statistical comparison to YAC128 mice injected with AAV9-CAG-CBE-mRosa26 (n = 7). **(c)** (Left) Sashimi plot depicting the splicing outcomes in the striatum of 12-month-old YAC128 mice injected with 1 x 10^11^ VGs of AAV9-CAG- SpRY-ABE-HTT-3 (n = 5) or AAV9-CAG-SpRY-ABE-mRosa26 (n = 7). (Right) Summary of HTT exon 13 skipping rates, as determined by NGS. **(d)** (Left) Western blot for human HTT protein products in the striatum of 12-month-old YAC128 mice injected with 1 x 10^11^ VGs of AAV9-CAG- SpRY-ABE-HTT-3 (n = 3) or AAV9-CAG-SpRY-ABE-mRosa26 (n = 3). (Right) Quantification of relative protein. Values are normalized to GAPDH. **(e, f)** Normalized **(e)** striatum and **(f)** cortex volumes of 12-month-old YAC128 mice injected with 1 x 10^11^ VGs of AAV9-CAG-SpRY-ABE- HTT-3 (n = 4) or AAV9-CAG-SpRY-ABE-mRosa26 (n = 7). Data are normalized to age-matched wild-type FVB/NJ mice (n = 5). **(g)** Representative immunofluorescence staining of mHTT inclusions in the striatum of 12-month-old YAC128 mice injected with 1 x 10^11^ VGs of AAV9-CAG- SpRY-ABE-HTT-3 (2 sections analyzed per biological replicate; n = 5) or AAV9-CAG-SpRY-ABE- mRosa26 (2 sections analyzed per biological replicate; n = 6). Scale bar, 52 µm. **(h, i, j, k)** Quantification of **(h)** the percentage of cells with mHTT inclusions, **(i)** the percentage of cells with ≥15 mHTT inclusions, **(j)** the percentage of cells with mHTT inclusions per unit area, and **(k)** the percentage of cells with ≥15 mHTT inclusions per unit area. All data are normalized to YAC128 mice injected AAV9-CAG-SpRY-ABE-mRosa26. Values are means and error bars indicate S.D. *P < 0.05, **P < 0.01, *** P < 0.001, ****P < 0.0001. Data for **(b)**, **(d, e)**, and **(h, i, j, k)** compared using a one-tailed unpaired t-test. Data for **(c)** compared using a one-way ANOVA with Tukey’s post-hoc analysis.

Using NGS, we analyzed editing outcomes in bulk striatal tissue at two time-points: one-month and four-months post-injection. From this analysis, we observed a dose- and time-dependent increase in A > G editing at the exon 13 SA (**Fig. S9**), with editing rates of ∼0.6%, ∼1.4%, and ∼4.5% observed at one-month post-injection for mice injected with 1 x 10^9^, 1 x 10^10^, and 1 x 10^11^ VGs per hemisphere, respectively (P < 0.001; **Fig. S9**), and editing rates of ∼1.1%, ∼2.7%, and ∼8.1% observed at four-month post-injection for mice injected with 1 x 10^9^, 1 x 10^10^, and 1 x 10^11^ VGs per hemisphere, respectively (P < 0.01; **Fig. S9**). Because the editing rate for SpRY-ABE-HTT-3 was approximately four-fold higher when delivered at 1 x 10^11^ VGs versus 1 x 10^10^ VGs (P < 0.01), we elected to use the higher dosage for all subsequent studies.

Following this dose-escalation, we used NGS to analyze editing outcomes in bulk striatal tissue from 12-month-old YAC128 mice injected with 1 x 10^11^ VGs of the split-intein ABE, which revealed the target edit in ∼16% of all reads (**Fig. 5b**, P < 0.0001). Notably, ∼15% and ∼3% of all reads were also found to carry the above-described bystander edit +7-bps and +9-bps downstream of the target SA, respectively (**Fig. 5b**). Using NGS, we also measured the frequency of exon skipping on PCR products from cDNA prepared from bulk striatal tissue, observing an overall skipping rate of ∼9.1% (P < 0.0001; **Fig. 5c**), with ∼4.9% of all transcripts found to lack all of exon 13 (P < 0.0001) and ∼4.2% of transcripts found to lack only the first 39-bps of exon 13 (P < 0.0001; **Fig. 5c**), which was altogether consistent with our findings from HEK293T cells.

Using the anti-HTT antibody clone 5HU-1H6, which recognize residues 115-129 of the human protein, we determined the effects of SpRY-ABE-HTT-3 on the HTT protein by western blot. Within the striatum, we found that 12-month-old YAC128 mice treated with SpRY-ABE-HTT- 3 had ∼60% less N-terminal HTT protein (P = 0.013; **Fig. 5d**) and ∼28% less FL HTT protein (P = 0.07; **Fig. 5d**) compared to the controls.

To determine if the split-intein ABE improved HD-related deficits, we used MRI to monitor volumetric changes in the striatum and cortex of YAC128 mice. While 12-month-old YAC128 mice injected with the mRosa26-targeting ABE had a ∼22% and ∼20% decrease in their striatal and cortical volumes, respectively, compared to the wild-type littermates (**Fig. 5e, f**), YAC128 mice treated with the split-intein ABE had only a ∼3% and ∼2% reduction in these respective volumes (P < 0.05 for both; **Fig. 5e, f**) compared to wild-type mice. We further quantified the density of neurons in the striatum and cortex of 12-month-old YAC128 mice (**Fig. S10a**), measuring that animals injected with SpRY-ABE-HTT-3 had ∼22% and ∼19% more NeuN^+^ cells in the striatum (P = 0.006; **Fig. S10b**) and cortex (P = 0.015; **Fig. S10c**) compared to the age-matching controls injected with an mRosa26-targeting ABE.

Finally, using the EM48 antibody, which recognizes aggregated forms of the mHTT protein, we conducted an immunofluorescence-based analysis to determine if the ABE reduced mHTT inclusions in the striatum of 12-month-old YAC128 mice. Compared to controls, mice treated by base editing had a ∼63% decrease in the percentage of cells with detectable mHTT inclusions (P < 0.0001; **Fig. 5h**), an ∼82% decrease in the number of inclusions per cell (P < 0.0001, **Fig. 5i**), and a ∼56% decrease in the number of cells with inclusions per area unit (P < 0.001; **Fig. 5j**). Additionally, among cells with detectable mHTT inclusions, we measured a ∼78% decrease in the total number of cells with >15 inclusions for mice treated by base editing (P < 0.0001; **Fig. 5k**).

In conclusion, we have developed CRISPR base editing platforms capable of disrupting the SA for HTT exon 13, which we show can induce its skipping to produce HTT protein isoforms resistant to proteolysis by caspase-6. These platforms were delivered by AAV to the striatum of a transgenic rodent model, where their editing decreased the accumulation of mHTT inclusions and led to the preservation of striatal and cortical volumes. Our results demonstrate the potential of base editing for treatment of HD.

## DISCUSSION

In this work, we describe an approach to treat HD using CRISPR base editing, which we find can improve two hallmarks of the disease: the accumulation of mHTT aggregates in the brain, and striatal and cortical volume loss. Given the critical role that the mHTT protein plays in HD [9, 10] and that its genetic depletion from cells can reverse features of HD [73–75], both antisense oligonucleotides (ASOs) and RNA interference (RNAi) have been used to lower the mHTT protein [75–80], with tominersen, an ASO developed by Ionis Therapeutics, and AMT-130, a microRNA-based gene therapy developed by uniQure, in clinical trials. However, tominersen and AMT-130 are non-allele-specific and thus target both the mutant and wild-type HTT transcripts, which has raised concerns about tolerability, as HTT has been found to play an important role in numerous physiological processes [81, 82]. As an alternative, allele-specific approaches that target single nucleotide polymorphisms (SNPs) associated with the mHTT allele have been explored [83–85], including several ASOs developed by Wave Sciences that have advanced to clinical testing (SELECT-HD; NCT05032196). However, these strategies are limited not just by the number of targetable SNPs available within the HD patient population, but also the financial and technical demands associated with individualized therapies. More broadly, these efforts suffer from other limitations, including the transient lifecycle of ASOs, which require a lifetime of administrations that could pose a burden on patients and lead to periods of diminished activity. Further, tominersen, the non-allele-specific ASO for HTT developed by Ionis, was found to trigger a dose-dependent ventricular enlargement, a safety concern that could lead to life-threatening side effects [86]. Conversely, while miRNAs and shRNAs can be expressed for an extended period of time from a viral vector to continuously engage with a target mRNA, these modalities rely on endogenous RNA processing pathways, whose saturation carries risk for dysregulating certain cellular functions [87]. Thus, there is a major need for new approaches that not only have the ability to permanently lower the mHTT protein but can do so without fully depleting the reservoir of the wild-type protein by a manner applicable to the entire HD patient population.

To overcome the limitations of existing gene therapies for HD, we explored the potential for CRISPR base editing technology to skip exon 13 of the HTT gene, both in cell culture models and a transgenic rodent model of HD. While HTT has been targeted and disrupted using CRISPR- Cas9 nucleases [88, 89], we used base editors in this study given that: (1) they feature more homogeneous and predictable editing outcomes compared to traditional nucleases and (2) they possess a more favorable safety profile. In particular, as base editors do not rely on DSBs nor NHEJ to initiate editing, they carry a reduced risk for inducing various genotoxic effects, including chromosomal deletions, translocations and chromothripsis [90–93].

Critically, the base editing approach used here to disrupt the SA for exon 13 produced two major transcript products: one that contained a deletion of all of exon 13, which led to a premature stop codon that decreased HTT expression, and another that contained a deletion of the first 39- bps of exon 13, which removed four of the amino acids in the caspase-6 cleavage site. While the function of caspase-6 in HD has been debated [94], it’s nonetheless been demonstrated that modifying its cleavage site in the mHTT protein can improve HD-related deficits in YAC128 mice, preserving striatal and cortical volumes for up to 12 months of age [12, 45, 95], a finding consistent with our results. Importantly however, our study did not address if the novel HTT isoform generated by our strategy is deficient at any normal physiological role. Additional studies will be needed to address this question. Nonetheless, unlike small molecule- [96] or antibody-based [97] approaches aimed at blocking the proteolysis of mHTT by caspase-6, our approach disrupts the protease cleavage site(s) itself and may thus carry a reduced risk for inducing adverse effects compared to those that pharmacologically inhibit caspase-6, an enzyme that contributes to many other critical physiological functions [98–101].

In addition to targeting exon 13 of the HTT gene, we also identified base editing systems capable of disrupting splicing elements in HTT exon 12. Though not extensively characterized in our study, HTT exon 12 represents a viable potential target for skipping, as evidenced by prior work that found that producing HTT protein isoforms that lacked the final 45 amino acids encoded by this exon reduced the formation of the N-terminal HTT protein fragments and improved deficits in a rodent model of HD [15, 102].

Previous studies have shown that deleting Hdh, the murine analog of HTT, is embryonic lethal mutation in mice and that its postnatal silencing can reduce lifespan, induce progressive motor impairments and cause a neuropathology, cautioning against the widespread, prolonged non-selective suppression of HTT in humans [81, 82]. Interestingly, heterozygous Hdh (+/−) mice do not manifest a Hdh-deficient phenotype, suggesting that the partial suppression of HTT may be comparatively safer [103]. Given our exon skipping-based approach only partially reduced the FL HTT protein, we note that our strategy may offer a potentially safer alternative to the non-allele-specific approaches developed to date for silencing mHTT. Of note, a limitation to this study is that YAC128 mice still possess the endogenous mouse Hdh gene. This functional redundancy can obscure the collateral effects of partially or completing depleting the wild-type HTT protein by a non-allele-specific strategy. In order to more comprehensively evaluate the tolerability of the approach described here, it will be critical to conduct studies in additional animal models, such as humanized Hu128/21 mice [104], which carry both a mutant and healthy copy of the human HTT gene in lieu of both copies of the mouse Hdh gene.

Neuropathological investigations suggest that HD is marked by a distinctive pattern of neuronal loss and subsequent brain atrophy [105–108] in the striatum and cortex [109–111]. Interestingly, despite observing DNA editing and exon skipping rates lower than 20% in bulk tissue from the striatum, we observed a robust preservation of both the striatum and cortex in YAC128 mice, as animals treated by our most effective platform, SpRY-ABE-HTT-3, were found to have only a ∼3% decrease in striatal volume at 12 months of age. We observed a similar beneficial effect for the mHTT inclusions, finding that animals treated by our approach had ∼63% less inclusions at 12 months of age. This could be explained in part by the observation that the mHTT protein is able to spread via cell-to-cell [112] and through the late endosomal secretory pathway [113], which can lead to the accumulation of mHTT aggregates in neighboring cells and, in some cases, trigger the aggregation of the wild-type HTT protein [114]. We hypothesize that modulating the abundance of the mHTT protein at a relatively early-stage of disease may have inhibited a feed-forward control mechanism, which resulted in an effect that is not entirely commensurate with the degree of editing measured in our bulk tissue analyses.

Last, OT effects are a crucial point to consider for any strategy that relies on a DNA editing platform [115, 116]. Our analysis into these effects, which measured editing at computationally predicted OT sites, found no evidence for any unintended mutations. However, further analyses utilizing emerging unbiased-based methods capable of detecting OT editing on a genome-wide scale, such as EndoV-seq [117] will be necessary to more precisely characterize the specificity of our approach.

In summary, we demonstrate that exon skipping with CRISPR base editing can be used to create new HTT protein isoforms resistant to proteolysis by caspase-6, which we found mitigated the development of HD-related deficits in a transgenic rodent model of the disease. Our results demonstrate the broad potential of CRISPR base technology for HD and illustrate promise for exon skipping to permanently produce forms of the mHTT protein with decreased toxicity.

## METHODS

### Plasmids

Plasmids encoding BE3 (Addgene #73021) [16], xCas9(3.7)-BE3 (Addgene #108380)) [31], xCas9(3.7)-ABE7.10 (Addgene #108382)[31], SaKKH-BE3 (Addgene #85170) [32], SpRY-CBE4max (Addgene #139999) [71] and ABE8e (Addgene #138489) [70] were obtained from Addgene. The plasmid encoding ABE7.10 was previously described [118]. To construct SpRY- ABE8e, Gibson assembly was used to insert a gBlock (IDT) encoding the SpRY SpCas9 mutations (D1135L, S1136W, G1218K, E1219Q, N1317R, A1322R, R1333P, R1335Q, and T1337R) into the EcoRV and AgeI restriction sites of pABE8e [70].

To construct the split-intein plasmid pAAV-CAG-N-SpCas9-CBE, Gibson assembly was used to insert a gBlock containing a CAG promoter and the N-terminal domain of the split-intein CBE (i.e., the V5 epitope tag, the rAPOBEC1 domain with a 16 residue linker [16], residues 1- 712 of the SpCas9 protein, and the N-intein from the DnaB protein of *R. marinus*) with the U6 promoter and sgRNA into the XbaI and NotI restriction sites of pX602 (Addgene #61593) by Gibson Assembly. To construct pAAV-CAG-C-SpCas9-CBE, the C-terminal domain of split-intein CBE (i.e., the C-intein fragment from the DnaB protein, residues 713-1,371 of SpCas9, the uracil glycosylase inhibitor, three repeats of the HA epitope tag, and an SV40 nuclear localization signal [NLS] sequence) with the U6 promoter and sgRNA were inserted into the XbaI and NotI restriction sites of pX602 by Gibson assembly.

To construct the split-intein pAAV-CAG-N-SpRY-ABE8e, Gibson assembly was used to insert a gBlock encoding the N-terminal domain of the split-intein ABE (i.e., the V5 epitope tag, an NLS, the ABE8e deaminase domain and a 32 amino acid linker [17] was inserted into the AgeI and SalI restriction sites of the plasmid pAAV-CAG-N-SpCas9-CBE, which was described above. To construct pAAV-CAG-C-SpRY-ABE8e, Gibson assembly was used to insert a gBlock containing the C-terminal domain of the split-intein ABE (i.e., the DnaB protein of *R. marinus* and residues 713-1,088 of SpCas9) into the AgeI and SalI restriction sites of pAAV-CAG-N-SpCas9- CBE. The ensuing plasmid was then used as a backbone to clone a gBlock containing residues 1,089-1,371 of SpCas9 with the SpRY mutations (D1135L, S1136W, G1218K, E1219Q, N1317R, A1322R, R1333P, R1335Q, and T1337R) with a 10 amino acid linker into the SalI and BamHI restriction sites. All amino acid sequences are provided in **Table S1.**

To construct the sgRNA expression vectors, oligonucleotides encoding the sgRNA targeting sequences were acquired from IDT (**Table S2**) and hybridized, phosphorylated and ligated into BbsI and BsaI restriction sites of pSP-gRNA (Addgene #47108) and pAAV-CAG-split-intein-CBE/ABE, respectively [119]. All plasmid sequences were verified by Sanger sequencing.

### Cell culture and transfection

HEK293T cells were obtained from the American Type Culture Collection (ATCC) and maintained in Dulbecco’s Modified Eagle Medium (DMEM) supplemented with 10% fetal bovine serum (FBS) and 1% penicillin/streptomycin at 37°C with 5% CO2. HEK293T cells were transfected in 24-well plates using Lipofectamine 2000 (Thermo Fisher Scientific) according to the manufacturer’s instructions, with 500 ng of base editor-encoding plasmid and 500 ng of sgRNA-encoding plasmid used per well. For experiments using the split-intein versions of a base editor, 500 ng each of the N- and C-terminal plasmids was used.

### Clonal cell line generation

HEK293T cells were transfected with 500 ng of plasmid encoding pCMV-N-CBE-HTT-4 and 500 ng pCMV-C-CBE-HTT-4 using Lipofectamine 2000. At 24 hr post-transfection, cells were re-plated on a 10-cm dish at a 1:100 ratio and grown as colonies from single cells, with Sanger sequencing used to confirm the target edit in PCR amplicons generated from genomic DNA from randomly selected clones, as described below.

### Analysis of DNA editing

Genomic DNA was isolated using a DNeasy Blood and Tissue Kit (Qiagen) with PCR amplification performed using a KAPA2G Robust PCR Kit (KAPA Biosystems) using 20-100 ng of DNA, Buffer A (1x), Enhancer (1x), dNTPs (0.2 mM), forward primer (0.5 µM), reverse primer (0.5 µM), KAPA2G Robust DNA Polymerase (0.5 U) and water (up to 25 µL). Cycling parameters were used as recommended by the manufacturer.

Sanger sequencing of the PCR amplicons was performed by the W.M. Keck Center for Comparative and Functional Genomics at the University of Illinois Urbana-Champaign. Base editing efficiencies were estimated by analyzing sequencing chromatograms using EditR [120] with the primer sequences provided in **Table S3**.

### RT-PCR

RNA was harvested using the RNeasy Plus Mini Kit (Qiagen) according to manufacturer’s instructions. cDNA synthesis was performed using the qScript cDNA Synthesis Kit (Quanta Biosciences) with 1 µg of RNA. PCR was then performed using the KAPA2G Robust PCR Kit using 25 ng of cDNA, Buffer A (1x), Enhancer (1x), dNTPs (0.2 mM), forward primer (0.5 µM), reverse primer (0.5 µM), KAPA2G Robust DNA Polymerase (0.5 U) and water (up to 25 µL). Cycling parameters were used as recommended by the manufacturer.

PCR products were visualized using a 2% agarose gel stained with ethidium bromide and imaged using a ChemiDoc-It2 (UVP). The sequences for the primers used for each target are provided in **Table S3**.

### Densitometry analysis

Exon kipping efficiencies were determined by densitometry analysis. The PCR products obtained from RT-PCR were analyzed by agarose gel electrophoresis, with the intensity of the bands measured using ImageJ. After subtracting background, band intensity was calculated using the following formula: % exon skipping = (skipped band intensity) / (non-skipped band intensity + skipped band intensity), where band intensity is the sum of each pixel grayscale value within the selected area of the band.

### qPCR

qPCR was performed using the SsoFast EvaGreen Supermix (Bio-Rad) according to the manufacturer recommendations. 50 ng of cDNA and 500 nM of each qPCR primer were used per sample, with GAPDH used as the housekeeping gene. Thermocycling and cycle threshold (CT) measurements were conducted using the CFX96 Touch Real-Time PCR Detection System (Bio-Rad). Standard and melt curve analyses were performed to validate qPCR primers and cycling parameters (**Fig. S11**). Relative gene expression was determined using the delta-delta CT method [121]. All primers used for qPCR are provided in **Table S3**.

### Western blot

Cells were lysed using radioimmunoprecipitation assay (RIPA) buffer (10 mM Tris-HCl pH 8.0, 140 mM NaCl, 1 mM EDTA, 1% Triton X-100, 0.1% SDS, 0.5% sodium deoxycholate, 2.5% β-mercaptoethanol) or 100 μL 1x NuPAGE LDS Sample Buffer (Thermo Fisher Scientific) containing 2.5% (v/v) β-mercaptoethanol and subsequently boiled at 95°C for 5 min. Protein lysates were then electrophoresed on a NuPAGE Bis-Tris Protein Gel (4-12%) using NuPAGE SDS Running Buffer (Thermo Fisher Scientific) for 2 hr at 130 V before transfer to a nitrocellulose membrane in Towbin Buffer (20 mM Tris-HCl pH 8.3, 192 mM glycine, and 10% [v/v] methanol, 1% SDS) for 2.5 hr at 75 V in an ice chamber with constant agitation.

Membranes were then blocked for 1 hr with 0.5% (v/v) bovine serum albumin fraction V in Tris-buffered saline (TBS; 20 mM Tris-HCl pH 7.5, 150 mM NaCl) with 0.1%, Tween-20 (TBS-T).

Membranes were subsequently incubated overnight at 4°C with the following primary antibodies: rabbit anti-GAPDH (1:1,000, Cell Signaling Technology #2118) or rabbit anti-HTT, clone EPR5526 (1:1,000, Abcam #109115) in blocking solution. After overnight incubation, the membranes were washed three times with 1x TBS-T and incubated with HRP-conjugated goat anti-rabbit antibodies (1:2,500, Cell Signaling Technology #7074P2) in blocking solution for 1 hr at room temperature. Membranes were then washed three times with 1x TBS-T and developed using Clarity Western ECL Substrate (Bio-Rad) and visualized using an Odyssey Imager (LI-COR). Band intensity was quantitated using ImageJ and normalized to GAPDH.

### Next-generation sequencing

Genomic DNA and cDNA were isolated as described above. Amplicons for sequencing were generated by PCR using KAPA HiFi HotStart (Roche) using primers with overhangs compatible with Nextera XT indexing (IDT) (**Table S3**). Following validation by agarose gel electrophoresis, PCR products were isolated using AMPure XP PCR purification beads (Beckman Coulter). Indexed amplicons were generated using a Nextera XT DNA Library Prep Kit (Illumina) and subsequently quantified and pooled. Libraries were then sequenced using a MiSeq Nano Flow Cell for 251 cycles from each end using the MiSeq Reagent Kit v2 (500-cycles). FASTQ files were created and demultiplexed using bcl2fastq v2.17.1.14 Conversion Software (Illumina). All NGS was performed by the Roy J. Carver Biotechnology Center at the University of Illinois Urbana-Champaign (Urbana, IL).

Base editing rates were quantified using CRISPResso2 [122]. Reads with average Phred scores below 30 were removed, and remaining reads were aligned to the expected amplicon sequences.

Exon skipping rates were quantified using the STAR RNA-Seq aligner by Galaxy. Forward and reverse reads were combined and aligned to the human reference genome (GRCh38) using STAR. 2-pass mapping was then used for splice junction analysis with a MAPQ value of 60 for .bam files. Splice junctions were determined from the STAR SJ.out.tab files, where the percentage of a junction event was defined as the number of reads for the target junction divided by the total number of events at the specified junction. Sashimi plots for splice junctions were generated by the Integrative Genomics Viewer (IGV’s) using the .bam and .bai files produced by RNA-STAR. The Sashimi plots presented in this manuscript were traces of the images generated by IGV.

### AAV vector production

AAV vectors were manufactured as described previously [123]. HEK293T cells were seeded onto 15-cm plates and maintained in DMEM supplemented with 10% (v/v) FBS and 1% (v/v) penicillin/streptomycin. After 16 hr, cells were transfected with 65 μg total of pAAV-CAG- N/C-CBE-HTT-4 or pAAV-CAG-N/C-SpRY-ABE-HTT-3 alongside pAAV9 and pHelper in a 1:1:1 ratio using PEI MAX (pH 8). Cells were harvested 72 hr after transfection by manual dissociation using a cell scraper and centrifuged at 2,000 RPM for 5 min at room temperature. The supernatant was collected into a fresh tube and mixed with 40% PEG 8000 (Thermo Fisher Scientific) in a 4:1 ratio (v/v) and stored overnight at 4°C. Cell pellets were then resuspended in 2 mL of lysis buffer (50 mM Tris-HCl pH 8.0 and 150 mM NaCl) per plate. The following day, the solution was centrifuged at 3,000 RPM for 30 min at 4°C, with the resulting pellets resuspended in lysis buffer and subjected to three consecutive freeze-thaw cycles. Supernatants were subsequently treated with 0.5% Triton X-100 (Thermo Fisher Scientific) and 50 units/mL Benzonase (Millipore Sigma) and shaken at 37°C for 1 hr. The lysate was then centrifuged at 4,000 RPM for 15 min at room temperature. The resulting supernatant was overlaid onto an iodixanol density gradient using 15%, 25%, 40% and 60% Opti-Prep solution (Sigma-Aldrich) and the AAV vector was isolated by ultracentrifugation at 58,400 RPM for 2 hr at 18°C. This step was repeated using a second iodixanol density gradient using fractions of 30%, 40% and 60%. Following the second extraction, AAV vector was filter-dialyzed using an Amicon Ultra 100 kDa MWCO column (Millipore Sigma) with PBS containing 0.001% Tween-20. Following treatment with DNase I (Millipore Sigma), AAV vector titer was determined by qPCR, with the vector stored at −80°C.

### Stereotaxic injections

All animal procedures were approved by the Illinois Institutional Animal Care and Use Committee (IACUC) at the University of Illinois Urbana-Champaign and conducted in accordance with the National Institutes of Health (NIH) Guide for the Care and Use of Laboratory Animals.

YAC128 mice were generated by breeding female FVB/N mice with male YAC128 mice obtained from the Jackson Laboratory (Stock #004938). Genotype was determined by qPCR using transgene specific probes (TransnetYX).

For studies involving the CBE, two-month-old YAC128 mice were injected with 1 x 10^9^ VGs each of AAV9-CAG-N-CBE-sgRNA and AAV9-CAG-C-CBE-sgRNA in 2 μl of PBS with 0.001% Tween-20, per hemisphere. The coordinates for the injection were: anterior-posterior (AP) = 0.86 mm, medial-lateral (ML) = ±1.80 mm, and dorsal-ventral (DL) = −3.75 mm.

For studies involving the ABE, one-month old YAC128 mice were injected with 1 x 10^9^, 1 x 10^10^ and 1 x 10^11^ VGs of each AAV9-CAG-N-SpRY-ABE-sgRNA and AAV9-CAG-C-SpRY-ABE-sgRNA in 2 μl of PBS with 0.001% Tween-20, per hemisphere. The coordinates for each injection were: AP= 0.86 mm, MV = ±1.80 mm, and DL = −3.75 mm, −3.55 mm, and −3.35 mm.

All groups for each study were sex-balanced and litter-matched.

### Tissue harvesting and RNA purification

Mice were anesthetized using 3% isoflurane delivered by vaporizer in a closed chamber and transcardially perfused using PBS. The striatum and cortex were dissected and divided in two halves, with one half stored in RNAlater Stabilization Solution (Thermo Fisher Scientific) and the other flash frozen. DNA and RNA were isolated using the DNeasy Blood and Tissue Kit (Qiagen) and RNeasy Plus Mini Kit (Qiagen), respectively. Prior to purification, tissues were homogenized in PBS using a KIMBLE Dounce Tissue Grinder.

### Western blot for tissues

Protein was isolated from bulk striatal tissue using RIPA buffer supplemented 10 µL/mL of Halt Phosphatase Inhibitor Cocktail (Thermo Fisher Scientific 78420). Samples were then boiled at 95°C for 5 min and electrophoresed using a NuPAGE Bis-Tris Gel (4-12%) with NuPAGE SDS Running Buffer (Thermo Fisher Scientific) for 2 hr at 130 V, before a electrophoretic transfer to a nitrocellulose membrane in Towbin buffer with 0.5% SDS, for 2.5 hr at 75 V. Membranes were then blocked with 1% (v/v) non-fat dry milk in TBS-T for 1 hr and incubated overnight at 4°C with primary antibodies in blocking solution. The following primary antibodies were used: mouse anti-HTT antibody, clone 5HU-1H6 (1:1,000, Sigma Aldrich MAB5490) and rabbit anti-GAPDH (1:1,000, Cell Signaling Technology #2118). After the overnight incubation, membranes were washed three times with TBS-T and incubated with HRP-conjugated goat anti-mouse (1:1,000, Cell Signaling Technology #7056) or goat anti-rabbit (1:1,000, Cell Signaling Technology #7074P2) antibodies in blocking solution for 1 hr at room temperature. Membranes were washed three times with TBS-T, developed using Clarity Western ECL Substrate (Bio-Rad) and visualized using an Odyssey Imager (LI-COR). Band intensity was quantitated using ImageJ and normalized to GAPDH.

### Immunohistochemistry

Immunohistochemistry was performed as previously described [34]. Briefly, dissected striatum and cortex were fixed in 4% (v/v) paraformaldehyde (PFA) at 4°C and embedded in OCT for generation of 15 µm coronal sections using a CM3050 S cryostat (Leica).

For immunostaining, sections were washed three times with PBS for 15 min and incubated in blocking solution (PBS with 10% [v/v], goat serum [Abcam #7481] and 0.5% Triton X-100) for 2 hr at room temperature, at which point the sections were stained with primary antibodies in blocking solution overnight at 4°C. Afterwards, sections were washed three times with PBS and incubated with secondary antibodies for 1 hr at room temperature, before three final washes with PBS. Sections were then incubated with 1 μg/mL of DAPI (Sigma-Aldrich) for 10 min and mounted onto slides using VECTASHIELD HardSet Antifade Mounting Medium (Vector Laboratories). Sections were imaged using a Leica TCS SP8 confocal microscope (Beckman Institute Imaging Technology Microscopy Suite, University of Illinois Urbana-Champaign, Urbana, IL), with all images analyzed using ImageJ.

The following primary antibodies were used: rabbit anti-V5 (1:100, Cell Signaling Technology #13202), mouse anti-NeuN (1:400, Cell Signaling Technology #94403), rabbit anti- Iba1 (1:500; Wako Pure Chemicals Industries 019-19741), and mouse anti-GFAP (1:200; Cell Signaling Technology, #3670)).

The following secondary antibodies were used: goat anti-rabbit Alexa Fluor 555 (1:400, Cell Signaling Technology #4413) and goat anti-mouse Alexa Fluor 488 (1:400, Abcam #150113) and goat anti-chicken Alexa Fluor 647 (1:200, Abcam #150175).

### mHTT inclusion staining

Floating sections (30 µm) from the striatum and cortex were washed three times in 12- well plates with PBS and incubated with blocking solution (0.3% Triton X-100 and 1% goat serum in PBS) for 2 hr at room temperature. Sections were then incubated with the primary antibody EM48 (1:80, Millipore MAB5374) for 72 hr. Afterwards, sections were washed three times with PBS and incubated with the secondary antibody (goat anti-mouse Alexa Fluor 488, 1:400, Abcam 150113) for 2 hr at room temperature. Following a subsequent 10-min incubation with 1 μg/mL of DAPI (Sigma-Aldrich), floating sections were mounted onto microscope slides using ProLong Gold Antifade Mountant (Cell Signaling Technology #9071). Sections were then imaged using a Leica TCS SP8 confocal microscope (Beckman Institute Imaging Technology Microscopy Suite, University of Illinois Urbana-Champaign, Urbana, IL).

### MRI analysis

A 9.4T/20-cm horizontal bore animal MRI system with a cryoprobe and ParaVision v6.0.1 software (Bruker BioSpin) was used to analyze 12-month-old YAC128 and FVB/N mice. The scanner was equipped with a gradient coil set of 660 mT/m per axis and a 4570 T/m/s slew rate, allowing for high resolution imaging. Body temperature and respiratory function were monitored using a three-dimensional (3D) T2-weighted fast spin echo (FSE) sequence using a 1030 MRI-compatible small animal monitoring and gating system (SA Instruments). Mice were anesthetized with 2% isoflurane, which was delivered by a vaporizer.

Post-processing of MRI files in the form of DICOM images was conducted using 3D Slicer [124]. Briefly, anatomical images were first merged to generate an average of all anatomical images. This template was then segmented into different brain regions using the Allen Mouse Brain Atlas [125]. Transformations of the atlas were projected onto the images of the mouse brain and corresponding segmentation of brain anatomical structures were made. Striatal and cortical volumes were measured in mm^3^ and normalized to the volume of age-matched wild-type FVB/NJ mice.

## Statistical analysis

Statistical analysis was performed using GraphPad Prism v9.1 (GraphPad Software, Inc.). All experiments consisted of independent biological replicates. Groups were compared using an unpaired one-tailed t-test or a one-way ANOVA with Tukey’s post-hoc analysis.

## ACKNOWLEDGMENTS

We thank the DNA Services staff of the Roy J. Carver Biotechnology Center at the University of Illinois Urbana-Champaign, particularly Alvaro Hernandez and Chris Wright for support with NGS. We thank Shreyan Majumdar of the Biomedical Imaging Center at the Beckman Institute for Advanced Science and Technology for support with MRI. We thank the Imaging Technology Group at Beckman Institute for Advanced Science and Technology for support with confocal microscopy. We thank Richard Chen for helpful discussion. This work was supported primarily by the CHDI Foundation (A-16129). Additional support came from the National Institutes of Health (1U01NS122102, 1R01NS123556, 1R01GM141296, 1R01GM127497, 1R01GM131272), the Muscular Dystrophy Association (MDA602798), the American Heart Association (17SDG33650087), the Parkinson’s Disease Foundation (PF-IMP- 1950) and the Simons Foundation (887187). M.G. was supported by the GRFP program from the National Science Foundation (DGE 1746047). A.M. was supported by the National Institute of Biomedical Imaging and Bioengineering of the National Institutes of Health under Award Number T32EB019944. J.W was supported by the Northwestern University Clinical and Translational Science Institute (UL1TR001422). The content is solely the responsibility of the authors and does not necessarily represent the official views of the National Institutes of Health.

## AUTHOR CONTRIBUTIONS

M.G., T.G. and P.P conceived of the study; S.S., M.G., D.S., N.G., P.A., A.M., D.J., M.G.S., A.B., G.E., M.S., J.W., W.W., D.A., C.K.W.L designed and performed experiments and analyzed the data; S.S, T.G and P.P. wrote the manuscript with input from all authors.

