## Supplemental data for "In vivo CRISPR base editing for treatment of Huntington’s disease"

### SUPPLEMENTARY FIGURES

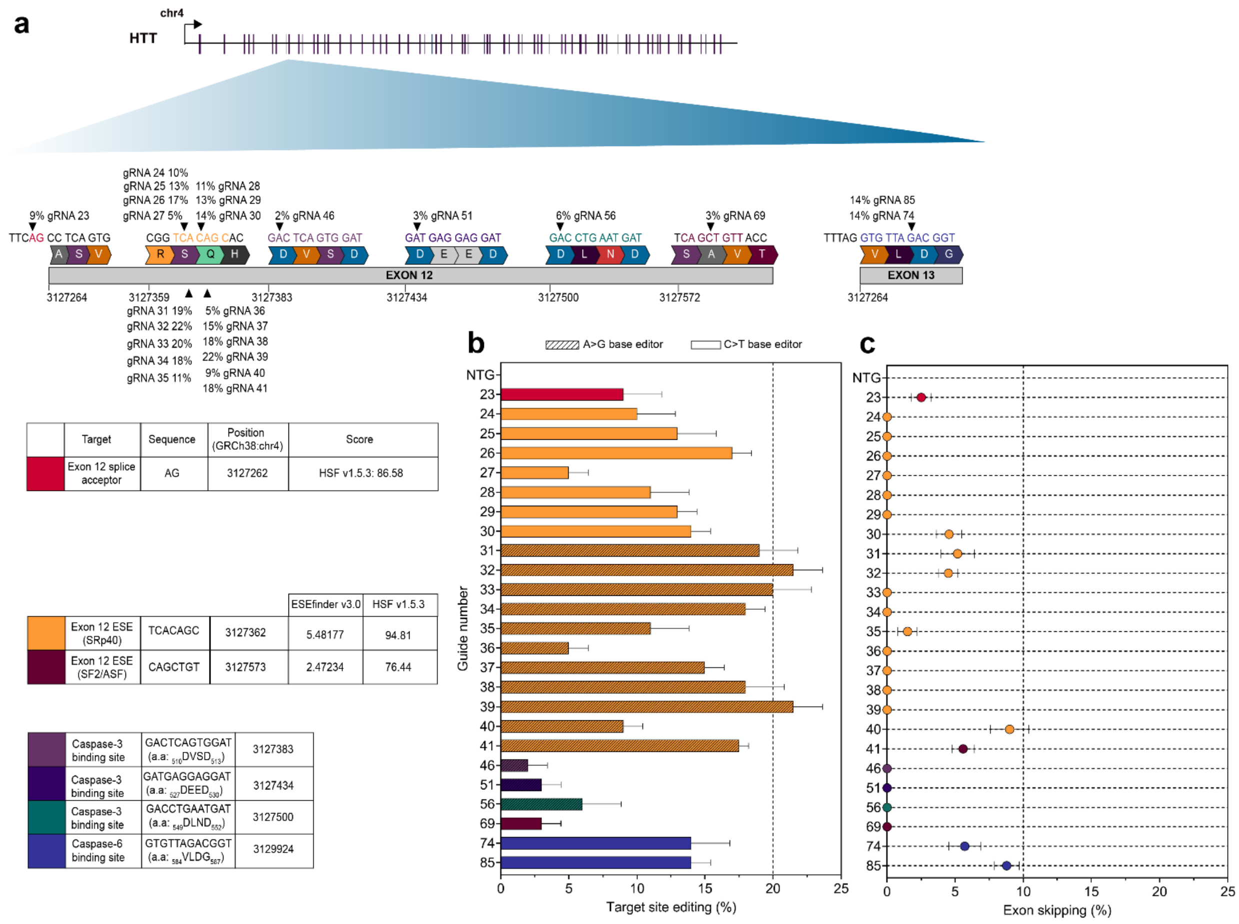


**Figure S1. CRISPR base editing of potential sites in HTT exon 12 and exon 13. (a)** Schematic representation of HTT and summary of target sites in HTT exon 12 and exon 13. **(b)** Genomic DNA editing rates in HEK293T cells with base editors targeting the splice acceptor of exon 12, 2 different splice enhancers within exon 12, 3 sequences encoding target sites for caspase-3 and a sequence encoding the target site for caspase-6 (n = 2). **(c)** Exon skipping rates following treatment with the base editors describe above measured by gel densitometry of RT-PCR products (n = 2). Values represent means and error bars represent S.D.

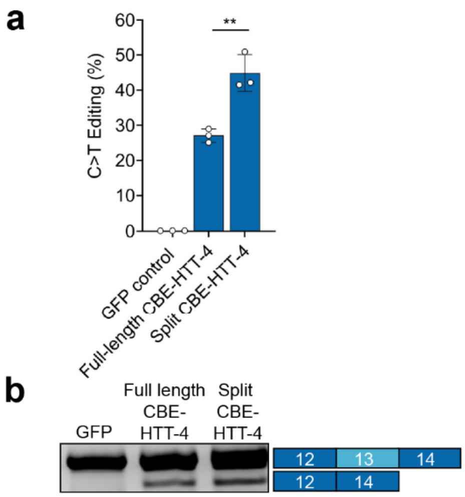


**Figure S2. Split CBE-HTT-4 outperforms its full-length counterpart when targeting the SA of HTT exon 13 *in vitro.*** **(a)** Base editing rates in genomic DNA in HEK293T cells at the SA of exon 13 following treatment with full-length or split CBE-HTT-4 quantified by NGS (n = 3). **(b)** Gel image of RT-PCR products detecting HTT exon 13 skipping following treatment with full-length or split CBE-HTT-4 (n = 3). Plasmid encoding GFP was used as a control in all experiments. Values represent means and error bars indicate S.D.; **, P < 0.01; Data compared using one-tailed unpaired t-test.

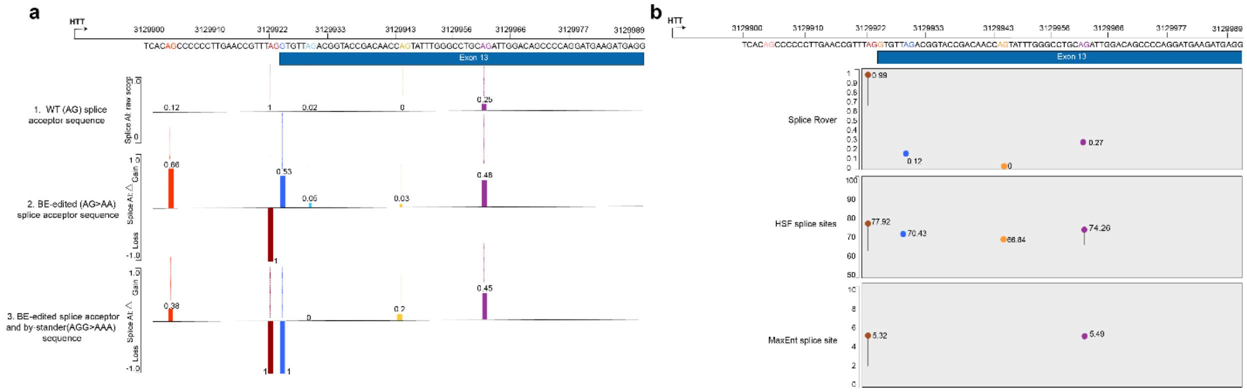


**Figure S3. Computational prediction of potential splice acceptor sites in HTT exon 13. (a)** Splice AI network predicts potential cryptic splice acceptor sites in HTT exon 13 in a sequence-specific context. Panel 1 shows the raw splice site scores at potential ‘AG’ sites in the wild-type HTT exon 13 sequence; panel 2 displays the predicted increase or decrease in score at splice acceptor sites when the consensus splice acceptor site “AG” is modified (AG > AA); panel 3 displays the predicted increase or decrease in scores at splice acceptor sites when consensus SA-editing is coupled by a bystander editing at +1-bp from the canonical SA site (AGG>AAA). Size and direction of bars represents the gain or loss in scores with respect to the raw splice site score of the wild type transcript in panel 1. **(b)** Strength of consensus splice acceptor and potential cryptic splice acceptor sites within HTT exon 13 as predicted by different algorithms including Splice Rover, Human Splicing Finder (HSF), and MaxEntScan.

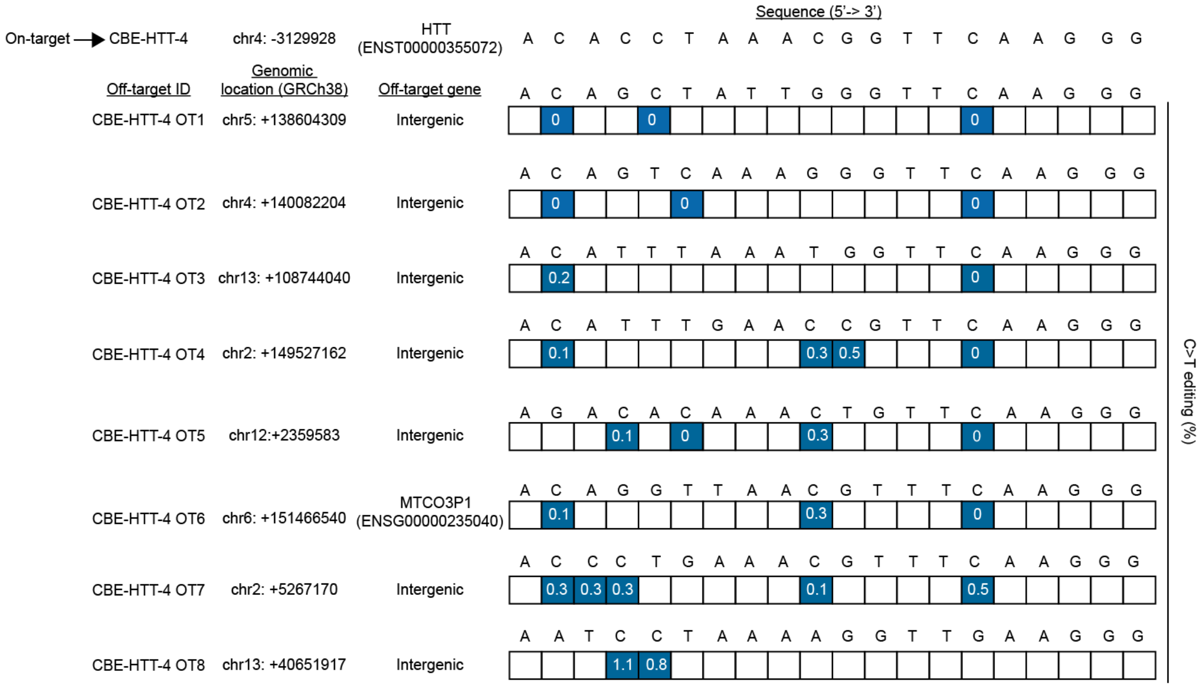


**Figure S4. CBE-HTT-4 did not edit computationally predicted off-target sites.** Chromosomal locations and gene IDs of the target sequence for CBE-HTT-4 and eight potential off-target sites identified using the specificity score described by Hsu et al. [1] is shown on the left. Tables describing the nucleotide frequencies for C > T editing, obtained by deep sequencing, for each off-target site is shown on the right. Frequencies are presented as the average percentage of edited reads (n = 3).

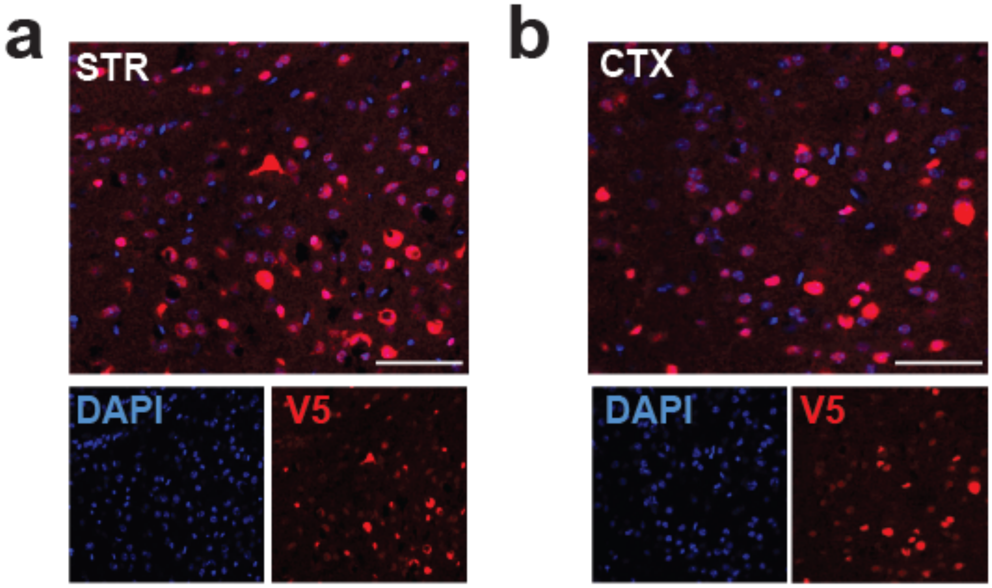


**Figure S5. AAV9-transduction of striatum and cortex in YAC128 mice.** Representative immunofluorescence images of **(a)** striatum (STR) and **(b)** cortex (CTX) in the brain of YAC128 mice injected with 1 x $10^9$ VGs of AAV9-CAG-CBE-HTT-4 per hemisphere. Scale bar, 50 µm. Staining for DAPI (blue) and anti-V5 antibodies (red).

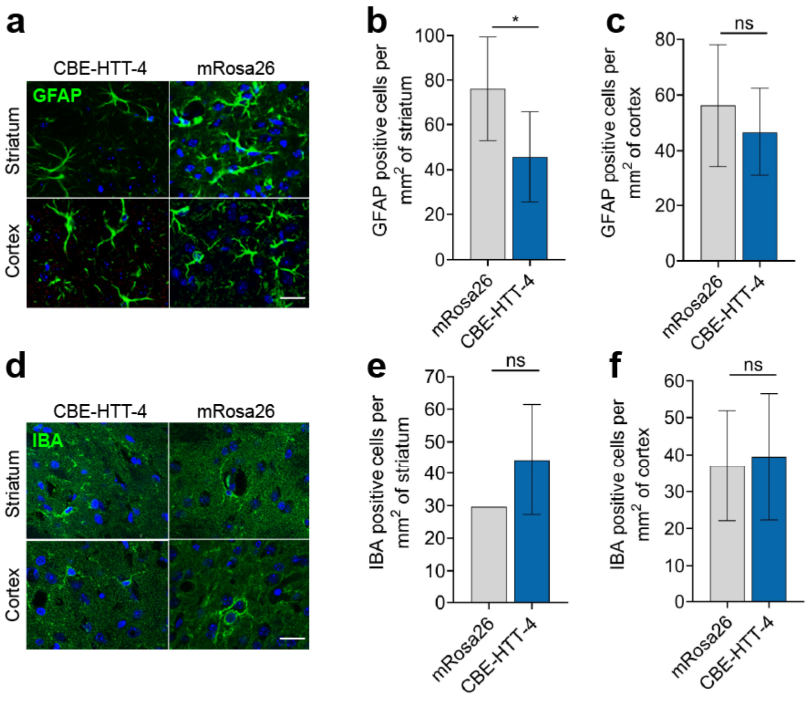


**Figure S6. Targeting HTT exon 13 SA with CBE-HTT-4 did not induce neuroinflammation in the striatum or cortex in YAC128 mice.** Representative immunofluorescence staining and corresponding quantification for **(a-c)** GFAP and **(d-f)** Iba1 in sections from the striatum and cortex of YAC128 mice injected with AAV9-CAG-CBE-HTT-4 or AAV9-CAG-CBE-mRosa26. Scale bar, 22.7 μm. Error bars indicate S.D. (n = 3). *, $P < 0.05$. Data for **(b-c), (e-f)**, compared using one-tailed unpaired t-test.

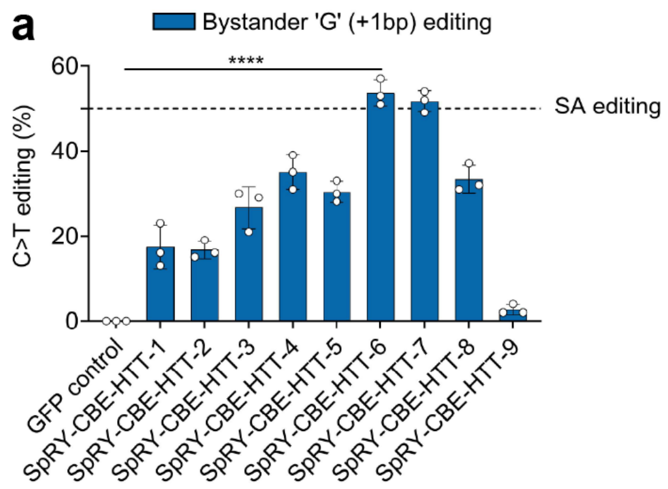


**Figure S7. Bystander editing induced by SpRY variants targeting the SA of HTT exon 13.** Genomic DNA editing rates measuring bystander edits using **(a)** SpRY-CBE4max or **(b)** SpRY-ABE8e in HEK293T cells (n=3). Values represent means and error bars indicate S.D. Data compared using one-tailed unpaired t-test.

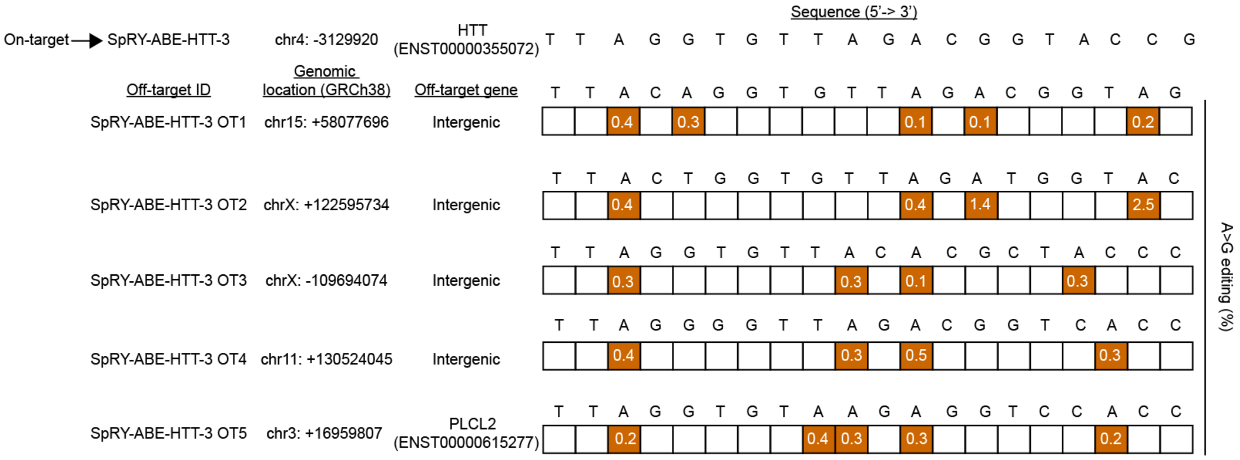


**Figure S8. SpRY-ABE-HTT-3 did not edit computationally predicted off-target sites.** Chromosomal locations and gene IDs of the target sequences for SpRY-ABE-HTT-3 and five potential off-target sites identified using Cas-OFF finder [2] are shown to the left. Tables describing the nucleotide frequencies for A > G editing, obtained by deep sequencing, for each off-target site is shown on the right. Frequencies are presented as the average percentage of edited reads (n = 3).

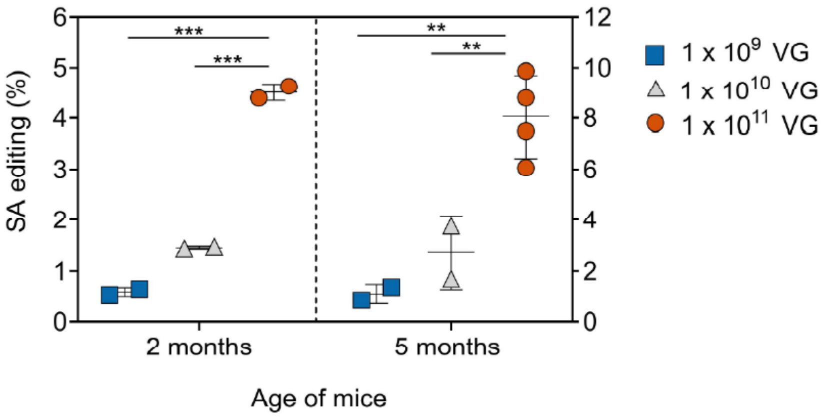


**Figure S9. AAV9 dose escalation study.** Genomic DNA editing rates of HTT exon 13 SA in YAC128 mice injected in the striatum with CAG-SpRY-ABE-HTT-3 packaged into AAV9. The doses used were 1 x $10^9$ VGs, 1 x $10^{10}$ VGs, and 1 x $10^{11}$ VGs. Measurements were performed at 2 months and 5 months of age. Values represent means and error bars indicate S.D. Data compared using one-tailed unpaired t-test.

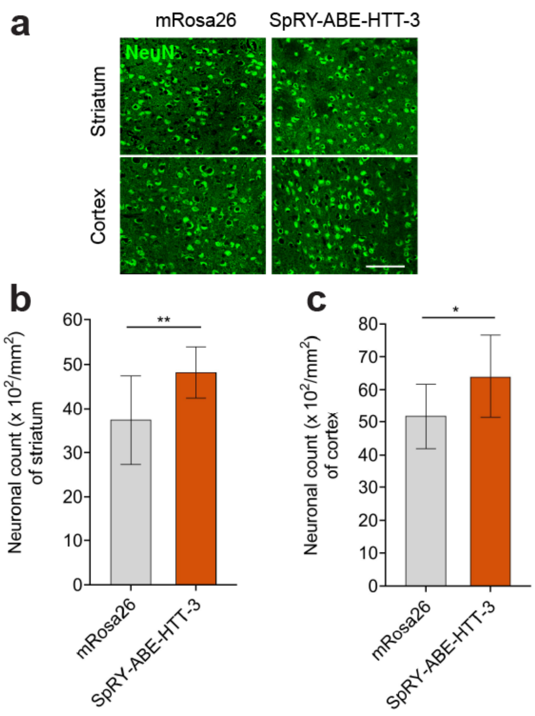


**Figure S10. Targeting HTT exon 13 SA site by AAV9-SpRY-ABE-HTT-3 affected neuronal count in the striatum and cortex regions of YAC128 HD mice. (a)** Representative immunofluorescence NeuN staining in sections from the striatum and cortex following injection of YAC128 mice with AAV9-CAG-SpRY-ABE-HTT-3 or AAV9-CAG-SpRY-ABE-mRosa26. Scale bar, 36.1 µm. **(b-c)** Quantification of NeuN-positive cells in the **(b)** striatum and **(c)** cortex in YAC128 mice with AAV9-CAG-SpRY-ABE-HTT-3 or AAV9-CAG-SpRY-ABE-mRosa26. Error bars indicate S.D. (n = 3). *, $P < 0.05$; **, $P < 0.01$. Data compared using one-tailed unpaired t-test.

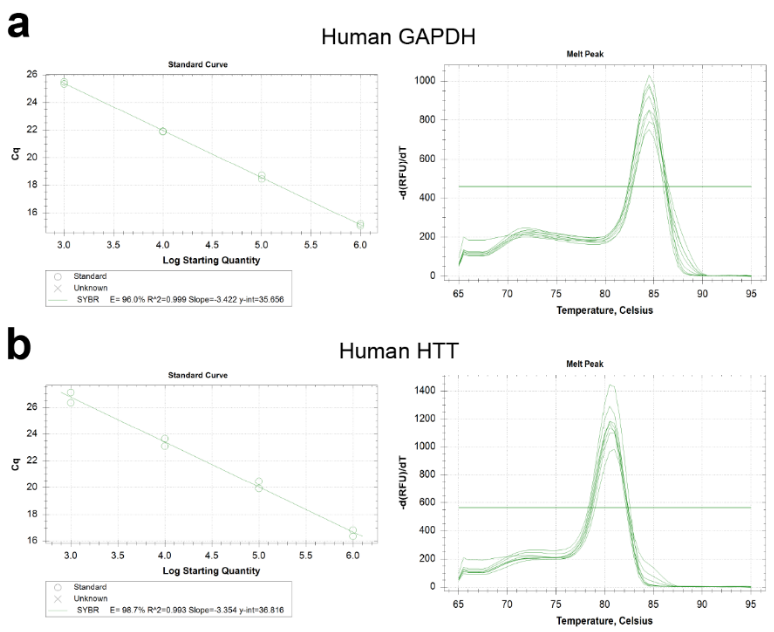


**Figure S11. qRT-PCR standard curves. (a)** Standard curve for qRT-PCR primers that bind to human GAPDH. **(b)** Standard curve for qRT-PCR primers that bind to human HTT.


1. Hsu PD, Scott DA, Weinstein JA, Ran FA, Konermann S, Agarwala V, Li Y, Fine EJ, Wu X, Shalem O, et al: **DNA targeting specificity of RNA-guided Cas9 nucleases.** *Nature Biotechnology* 2013, **31:**827-832.
2. Bae S, Park J, Kim J-S: **Cas-OFFinder: a fast and versatile algorithm that searches for potential off-target sites of Cas9 RNA-guided endonucleases.** *Bioinformatics* 2014, **30:**1473-1475.
